# Learning and actioning general principles of cancer cell drug sensitivity

**DOI:** 10.1101/2024.03.28.586783

**Authors:** Francesco Carli, Pierluigi Di Chiaro, Mariangela Morelli, Chakit Arora, Luisa Bisceglia, Natalia De Oliveira Rosa, Alice Cortesi, Sara Franceschi, Francesca Lessi, Anna Luisa Di Stefano, Orazio Santonocito, Francesco Pasqualetti, Paolo Aretini, Pasquale Miglionico, Giuseppe R. Diaferia, Fosca Giannotti, Pietro Liò, Miquel Duran-Frigola, Chiara Maria Mazzanti, Gioacchino Natoli, Francesco Raimondi

## Abstract

High-throughput screening platforms for the profiling of drug sensitivity of hundreds of cancer cell lines (CCLs) have generated large datasets that hold the potential to unlock targeted, anti-tumor therapies.

In this study, we leveraged these datasets to create predictive models of cancer cells drug sensitivity. To this aim we trained explainable machine learning algorithms by employing cell line transcriptomics to predict the growth inhibitory potential of drugs. We used large language models (LLMs) to expand descriptions of the mechanisms of action (MOA) for each drug starting from available annotations, which were matched to the semantically closest pathways from reference knowledge bases. By leveraging this AI-curated resource, and the interpretability of our model, we demonstrated that pathways enriched for genes crucial for prediction often matched known drug-MOAs and essential genes, suggesting that our models learned the molecular determinants of drug response. Furthermore, we demonstrated that by incorporating only LLM-curated genes associated with MOAs, we enhanced the predictive accuracy of our drug models.

To enhance translatability to a clinical setting, we employed a pipeline to align bulk RNAseq from CCLs, used for training the models, to those from patient samples, used for inference. We proved the effectiveness of our approach on TCGA samples, where patients’ best scoring drugs matched those prescribed for their cancer type. We further showed its usefulness by predicting and experimentally validating effective drugs for the patients of two highly lethal solid tumors, i.e. pancreatic cancer and glioblastoma.

In summary, our method facilitates the inference and interpretation of cancer cell line drug sensitivity and holds potential to effectively translate them into new cancer therapeutics.

**Highlights:**

-Interpretable drug-response prediction models were trained on large scale pharmacogenomics data sets (i.e. GDSC and PRISM).
-Large language models were used to enhance the curation of biological pathways associated to drugs MOA
-Unbiased interpretation of the models demonstrated learning of drug MOAs *and* gene essentiality
-Inference of TCGA cohort samples recovered mono- and combination cancer drug prescriptions and indicate potential repurposing candidates.
-Drug candidates predicted from bulk RNAseq samples of pancreatic cancer and glioblastoma were experimentally validated.

## Introduction

Landmark cancer genomic projects such as The Cancer Genome Atlas (TCGA) have provided an unprecedented, multimodal picture of the complex genetic and molecular landscape characterizing major tumor types (Chang et al., 2013). The possibility to define tumors at a molecular level is changing cancer treatment through the development of single- or combinatorial-targeted therapies for patients characterized by specific genetic makeups. Unfortunately, effective treatments are still lacking for many patients and the emergence of resistance further limits the clinical benefit of therapies.

In recent years, the availability of large-scale pharmacogenomics databases, including the Cancer Cell Line Encyclopedia (CCLE)(Barretina et al., 2012), the Genomics of Drug Sensitivity in Cancer v1 and v2 (GDSC)(Iorio et al., 2016), the Cancer Therapeutics Response Portal v2 (CTRPv2)(Rees et al., 2016) and the Profiling relative inhibition simultaneously in mixtures (PRISM)(Corsello et al., 2020), has fostered the development of personalized oncology strategies.

Initial landmark work by Iorio & coworkers highlighted the genomic alterations that sensitize to drugs as well as the contribution of different genome sequencing data types for drug sensitivity predictions (Iorio et al., 2016). Since then, several computational forecast methodologies have been proposed to leverage the information in CCLE and GDSC to predict cancer cell lines drug-sensitivity (reviewed in (J. Chen & Zhang, 2021; Firoozbakht et al., 2022; Xia et al., 2022) and best practices to develop predictive models have been issued (Sharifi-Noghabi et al., 2021). Models “primed” with prior knowledge have also been developed, that exploit representations of biological processes instead of individual genes as features to increase model performance and interpretability (Chawla et al., 2022; Ferraro et al., 2023; Kuenzi et al., 2020; Wang et al., 2019). However, a systematic investigation of whether drug sensitivity models are actually able to learn, in an unbiased fashion, the biological processes associated to drugs’ mechanism of action (MOA) are still missing from the literature. Moreover, to what extent models trained on expression data in cancer cell lines can be exploited to interpret patients’ bulk RNAseq data is still a largely unsolved issue, although earlier studies tried to tackle this fundamental point (Kuenzi et al., 2020; Ma et al., 2021).

In this study, to forecast drug sensitivity in cancer cell lines we developed a novel machine learning algorithm trained on large drug screening datasets, *CellHit*. We trained the models on GDSC and PRISM, as they represent the largest screening datasets of oncological and non-oncological drugs, respectively. We focused on interpretability and deployment of the predictive models, which are both critical to generate new testable hypotheses (Figure 1A). We employed our pipeline to infer the best-scoring drugs for the entire TCGA cohort based on patient transcriptomics profile, as well as on samples from PDAC and GBM patients, which were experimentally validated.

**Figure 1:**
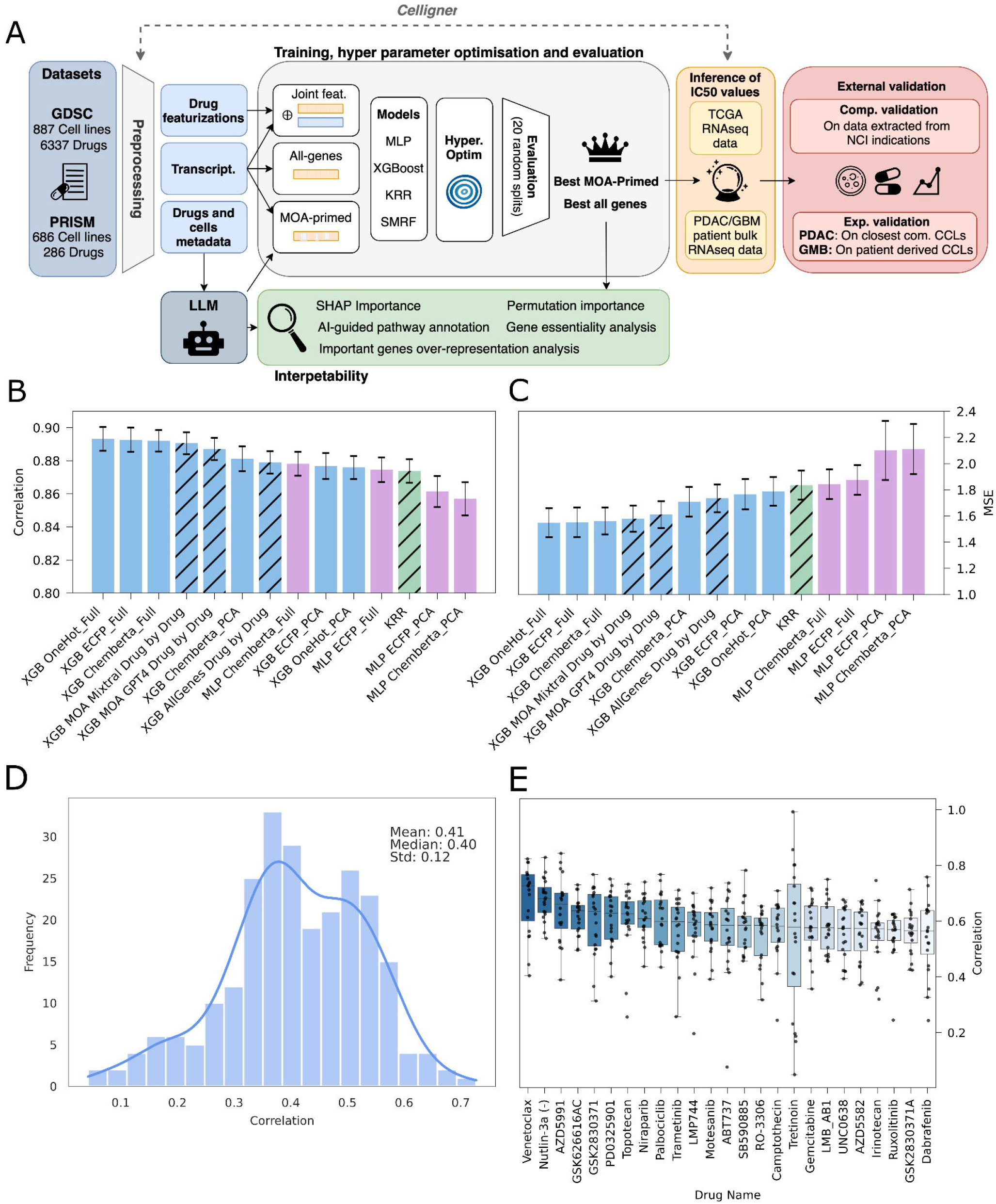
A) Schematic workflow of the pipeline from dataset acquisition and model training through to benchmarking and external validation. Leveraging large-scale transcriptomics datasets, several machine learning models are trained. A Large Language Model (LLM) assists the curation of drug Mechanisms of Action (MOA) enhancing model interpretability and facilitating feature selection. Three model types are trained: one using cell transcriptomics and drug features, another using only transcriptomics, and a third using a selected subset of transcriptomic data informed by pathways identified by the LLM. These models undergo benchmarking across 20 distinct train/validation/test splits. The best-performing model is then applied in inference on The Cancer Genome Atlas (TCGA) bulk RNA-seq data and on external patient datasets for pancreatic ductal adenocarcinoma (PDAC) and glioblastoma multiforme (GBM). Predictions are externally validated on TCGA data using National Cancer Institute (NCI) cancer drug indications to assess the recovery of known information, as well as experimentally on primary (GBM dataset) and commercial (PDAC dataset) cancer cell lines; B) Bar plot comparing the performance of different model architectures (MLP, XGBoost and literature baselines) and input feature representations (cell features and drug features) in terms of Pearson correlation with observed drug sensitivities. Different colors denote different learning algorithms (e.g light blue XGBoost and purple MLP). Etched bars highlight models using only transcriptomic data (no drug featurizations); C) Bar plot depicting Mean Squared Error (MSE) for the same models and features as in (B); D) Histogram of the distribution of Pearson correlation coefficients for drug-specific models using all genes, indicating the median, mean, and standard deviation; E) Box plots illustrating the variability in the distribution of Pearson correlation coefficients across 20 different random training/testing splits. Each box plot represents a specific model and displays the median correlation (central line), interquartile range (box edges), and variability outside the upper and lower quartiles (whiskers)

## Results

### An interpretable model for cell line drug response predictions with or without drug representations

We developed an interpretable machine learning framework for the prediction of drug sensitivity of cancer cell lines by using the latest version of the Genomics of Drug Sensitivity in Cancer (GDSC) dataset, which, after data pre-processing, provided the profiling of 286 unique drugs in 686 cell lines. We first implemented an integrated model by jointly considering representations of both drugs and cell lines to predict IC50 values as target variables (Figure 1A). We employed different strategies to numerically represent drugs chemical structures and cell line expression profiles to provide inputs to ML algorithms. As for cell lines, we first transformed the RNAseq data using Celligner (Warren et al., 2021) along with TCGA bulk RNAseq samples. This preprocessing step is required to align RNAseq data from cell lines to TCGA to apply our ML model on patient samples (see below). We employed drug and cellular representations as features to train a supervised regression model to predict IC50 data. We trained different algorithms and repeated the training procedure for each model by randomly splitting the training and testing sets 20 times (see Methods). We found that XGBoost, in combination with all-gene expression vectors and one-hot encodings of the molecules, achieved the best performance (Pearson correlation coefficient ρ=0.89, Mean Square Error MSE=1.55; Figure 1B,C), being able to outperform other architectures, such as MLP, as well as other competitive methods available from the literature (Chen & Zhang, 2021)(i.e. KRR, SRMF; Figure 1B,C; Supplementary Figure 1; Supplementary Table S1).

The fact that the simplest representation of the drug, i.e. one-hot encoding, achieved the best results, suggested that the model is not actually leveraging information from drugs’ structures, but rather using them as mere drug identifiers. Since our main goal was to learn the transcriptional programs responsible for drug responses for post-hoc interpretation, we generated drug-specific models by employing only gene expression as input features. We trained the models by repeating the same procedure used for the joint representation. Overall, aggregated IC50 predictions have correlations with experimental values almost as good as the joint model (ρ=0.88, MSE=1.73; Figure 1B,C; Supplementary Table 1), further confirming that drug representations didn’t significantly contribute to model performance. To assess the capability of our models to capture local variations in drug-cell line sensitivities, we measured whether the models not only approximated the mean IC50 value of a drug across various cell lines, but also reflected the distinct sensitivities exhibited by different cell lines. By evaluating performances on a held-out testing set, we obtained a median ρ of 0.40 out of 286 drug-specific models (Figure 1D). The best performance was achieved by Venetoclax (ρ=0.72), a small molecule regulating apoptosis by inhibiting BCL2 (Kapoor et al., 2020). Other top-performing models related to drugs involved in apoptosis regulation (the BCL2 inhibitor AZD5991), cell cycle regulation (the p53 pathway activators Nutlin-3a(-) and GSK2830371), inhibition of the DYRK3 kinase (GSK626616AC), ERK/MAPK signaling (PD0325901), DNA replication (the topoisomerase inhibitor Topotecan), genome integrity (the PARP inhibitor Niraparib) (Figure 1E). A total of 73 drug models (25%) had correlation performances greater than 0.5 (Supplementary Table S2). In summary, we developed reliable models for drug sensitivity based on just cell lines transcriptomics data.

### Models’ interpretation reveals convergence between important genes and known drug-targets

For each drug-specific model, we inspected the genes most important for prediction and checked whether they were either known targets, or components of pathways associated with the known MOA of that drug. We evaluated gene importance, defined as the contribution of individual genes to model predictiveness, in two ways. First, we inspected whether genes gave a positive or negative contribution to the final IC50 prediction via a game theory approach (i.e. SHAP). Second, we used an importance permutation method, randomly shuffling genes and evaluating the effect of the perturbation on test set metrics. We imposed a stringent criterion and considered only those genes found to be important with both approaches.

We found that 39% of the drug-specific models trained on GDSC identified the known target among the significant genes in at least one model out of 20 random splits (Supplementary Table S2). We used that fraction of model splits recovering the drug target to rank drug models. Remarkably, models for BCL2 inhibitors, such as Venetoclax, Navitoclax and ABT737, consistently recovered their target in the majority of the models trained on different splits (Figure 2A,B). Several other drug models recovered the corresponding targets in more than 50% of the splits (e.g. Gefitinib-EGFR, Nutlin-3a(-)-MDM2, or Linsitinib-IGF1R; Figure 2A). The recovery of the drug targets is significant, considering that 95.8% of the targets are found at or above the 90th percentiles of the background distributions of the frequencies of recovery of all genes across all splits for each drug (Figure 2A). Analysis of the importance scores, for example for the top performing model (Venetoclax), showed that higher values are attributed to the corresponding target, i.e. BCL2. This connection is seen both as a strong negative contribution (i.e. SHAP importance; Figure 2B teal) and in test metrics (i.e. Correlation delta; Figure 2B orange). By plotting for each cell line the target gene expression, Venetoclax’s experimental IC50s, as well as Venetoclax’s model SHAP values, it is possible to get a deeper insight into how the Venetoclax model leverages the expression levels of its corresponding target (i.e. BLC2) to successfully predict IC50 (Figure 2C). Indeed, cell lines having a higher expression of BCL2 are characterized by lower IC50 as well as strongly negative SHAP values (i.e. higher efficacy of Venetoclax as a BCL2 inhibitor) (Figure 2C).

**Figure 2:**
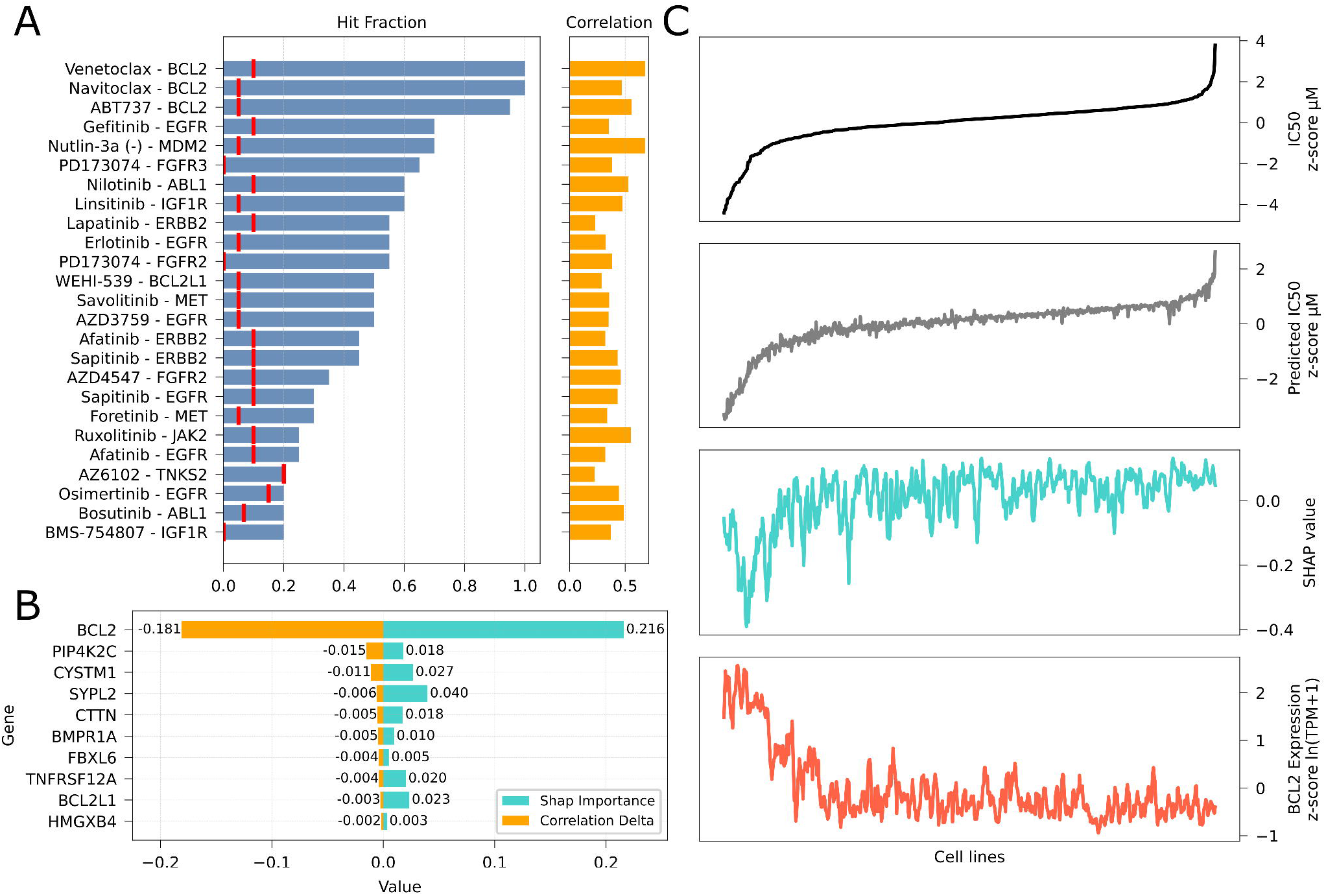
GDSC model interpretability. A) Target recovery for the top 25 ligand-target pairs. The length of the bars on the left indicates the fraction of recovery (# of times the drug-specific model identifies the putative gene as important out of 20 train/test splits). Red lines represent the 95th percentile of the Hit Fraction distribution across all genes for a given drug. The length of the bars on the right shows the median pearson correlation for each drug-specific model; B) SHAP (teal) and correlation delta (orange) importances for the Venetoclax drug. Permutation importance reflects the decrease in the model’s prediction accuracy when a feature’s values are shuffled, indicating its importance (greater drops signify higher importance). SHAP importance represents a feature’s contribution to the model’s prediction, with larger absolute values indicating greater importance; C) An integrated assessment of the Venetoclax model across various cell lines (X axis). The top plot (in black) shows the experimental IC50 z-scores, while the second plot (in gray) depicts the predicted IC50 values, providing a comparison of model performance against experimental data. The third and fourth plots (in teal and red) respectively represent the SHAP values and expression levels of BCL2. Overall, the figure shows how lower IC50 values (higher drug efficacy) are associated with higher BCL2 expression levels and correctly identified impact (negative SHAP value) of the gene on predicted IC50.

### MOA processes and gene dependencies are learnt by the drug sensitivity models

In addition to targets, we wanted to inspect whether the models learnt about MOA-related biological processes and pathways. However, the curated information of biological pathways associated to the action of a certain drug is only sporadically annotated in chemical bioactivity or biological pathway knowledge bases. We therefore set out to systematically curate the association of GDSC drugs to pathways in the reference knowledge-base, Reactome (Jassal et al., 2020). We first retrieved a total of 66 GDSC drugs that are annotated to Reactome pathways through Guide to Pharmacology/Pubchem (Harding et al., 2018) mappings. We then considered all the pathways containing the drug targets, finding matches for 233/287 (81%) unique GDSC drugs. To further extend the drug-MOAs’ pathway coverage, we leveraged open-source Large Language Models (LLMs), i.e. GPT4 and Mixtral Instruct, to generate detailed annotations of drug-related processes by inputting the generic name of the drug (or available synonyms) and available annotations from the GDSC compound information table. The resulting annotations informed a second prompt designed to identify semantically closest pathways from Reactome for each drug (Figure 3A; see Methods). Through this approach, we were able to retrieve detailed information about MOA and associated pathways for 253/287 (88%; Supplementary Table 3) GDSC drugs (Figure 3B, top). This procedure increased the coverage of annotated drugs, and it enabled the association of new pathways with the drug’s MOAs (Figure 3B, bottom). We finally combined the three sets of drug-pathway associations, for a total of 5662 instances, 253 unique drugs and 138 unique pathways (Supplementary Table 3).

**Figure 3:**
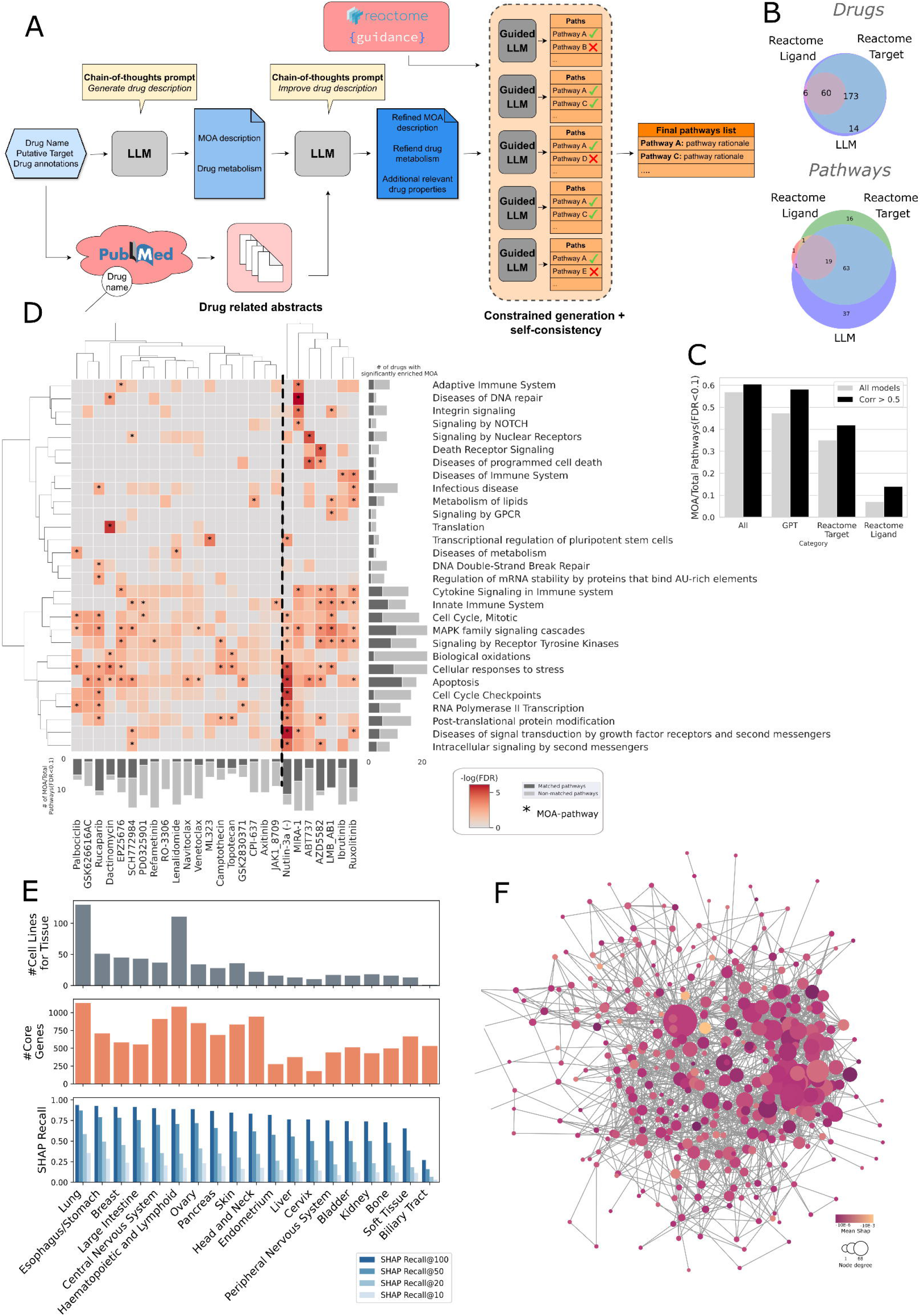
Analysis of Drug MOAs and Gene Essentiality via GDSC Models. A) Workflow depicting the use of a large language model (LLM) for generating drug MOAs and identifying semantically relevant pathways. Starting from the drug’s available metadata, an LLM is repeatedly tasked with specialized prompts to generate a drug textual description. In parallel, PubMed is queried programmatically with the drug name to retrieve abstracts related to the drug. The information is integrated in a final textual description. The obtained drug description is used by a “Guided” LLM to choose which are the Reactome pathways which are most likely to modulate drug efficacy. This last procedure is repeated 5 different times and only pathways selected at least two times are retained; B) Venn diagram showing the different drugs(top) and pathways (bottom) recovered using the LLM procedure as compared to pathway match based on drugs’ and target’s names; C) number of significantly enriched MOA-pathways obtained from different annotation criteria; D) Heatmap of significant MOA-pathways for various drug models, filtered by a correlation threshold >0.5. Drug names and involved pathways are labeled along the x-axis and y-axis, respectively. Starred squares highlight pathways linked to drugs via at least one annotation criterion. Adjacent bar plots show the count of significantly enriched elements per row/column in light gray, with those annotated by the pipeline in dark gray. A vertical dashed line highlights the presence of a group of drugs that most frequently recover pathways and known MOAs; E) tissue-wise statistics of number of cell lines (top), number of core essential genes (middle), recall of essential genes at different important genes (SHAP) stringencies (top k 10, 20, 50, 100); F) STRING PPI network of lung core essential genes recovered by SHAP importances. Nodes’ have diameters proportional to node degree and are colored according to SHAP values (the brighter the more important)

We performed pathway enrichment analysis on important genes to check whether known MOAs were identified. We found a total of 114 GDSC drug models with at least one significantly enriched pathway (FDR < 0.1). Out of these, 65 (57%) unique drugs have at least one MOA-pathway significantly enriched when considering the aggregated list of MOA-pathways (Figure 3C; Supplementary Table 4). When considering the different MOA-pathway sources, the LLM-derived one was the single list providing the highest recovery of enriched pathways (Figure 3C; Supplementary Table 4). The fraction of unique drugs with at least one, significantly enriched MOA-pathway slightly increased when considering models with a correlation greater than 0.5 (Figure 3C; Supplementary Table 4).

We then clustered pathway enrichment of the best-performing drug models (ρ > 0.5) having at least one significant MOA-pathway (Figure 3D). We found that certain processes, such as “Apoptosis”, “Cellular responses to stress”, “Biological Oxidations”, “MAPK family signaling cascades”, were widely enriched across drugs, often consistently with drugs’ MOAs, and clustered apart (Figure 3D). A second, larger cluster of pathways, including among others “Death Receptor Signaling” and “Integrin signaling”, was enriched for specific drugs and less frequently matched known MOA drugs (Figure 3D). Clustering also highlighted a group of drugs that most frequently recovered certain pathways related to cell cycle (Figure 3D, right cluster). Some of these had a prevalence of matched MOA among enriched pathways, such as LMB_AB1, a drug targeting adrenergic receptors ADRA1A and ADRB1. We also found many drugs with enriched pathways that did not match known drugs’ MOA (e.g. RO-3306, Figure 3D), indirectly suggesting potentially new MOAs.

We evaluated whether important genes also recovered information about gene dependencies of cancer cell lines from different tissues. In this respect, we retained drug-cell line instances yielding more significant predictions, ranked the top *k* most important genes based on SHAP values, and pooled them on the basis of the tissue of origin of the cell lines. We then evaluated the recall of the top *k* important genes to identify core essential genes from an updated dependency map across 27 cancer tissues (Pacini et al., 2024). Depending on how many top *k* important genes we considered, we were able to recover a variable number of gene dependencies for all the tissues. Remarkably, when aggregating the top 100 genes based on SHAP importance, we were able to identify core essential genes with a recall greater than 0.9 in several tissues (Figure 3E, Supplementary Table 5). These essential, prediction-important genes are typically found in protein-protein interaction networks (e.g. STRING network of important, essential genes in lung Figure 3F). By ranking genes based on their average SHAP importance across drug models, we found BCL2L1, YAP1 and CHKA as the top 3 most important genes (brighter nodes, Figure 3F; Supplementary Table 6). Overall, this analysis suggests that drug sensitivity is also realized through the modulation of critical network genes that are essential for cell survival and proliferation.

### Extending explainable drug sensitivity predictions to the PRISM dataset

We employed a similar strategy to train predictive models for 6337 drugs and 887 cell lines available in the PRISM database. We found that the majority of trained models were characterized by overall low performances (Figure 4A, blue curve and dots; median correlation=0.04), likely due to moderate and tissue-specific responses of cancer cell lines to the many non-oncological drugs present in PRISM. When considering only drug models characterized by dispersed experimental log-fold changes (LFCs) in their dataset, i.e. with InterQuartile Range (IQR) greater than 1, the average performance substantially increased (median ρ=0.24, red curve), yielding a total of 713 out of 6337 drug models. Based on this result, we decided to restrict the downstream analysis only to those models with ρ>0.2, as they represent most of those with IQR > 1 (Figure 4A). A cutoff of ρ>0.2 is moreover consistent with earlier modeling efforts on a previous release of the PRISM dataset (Corsello et al., 2020).

**Figure 4:**
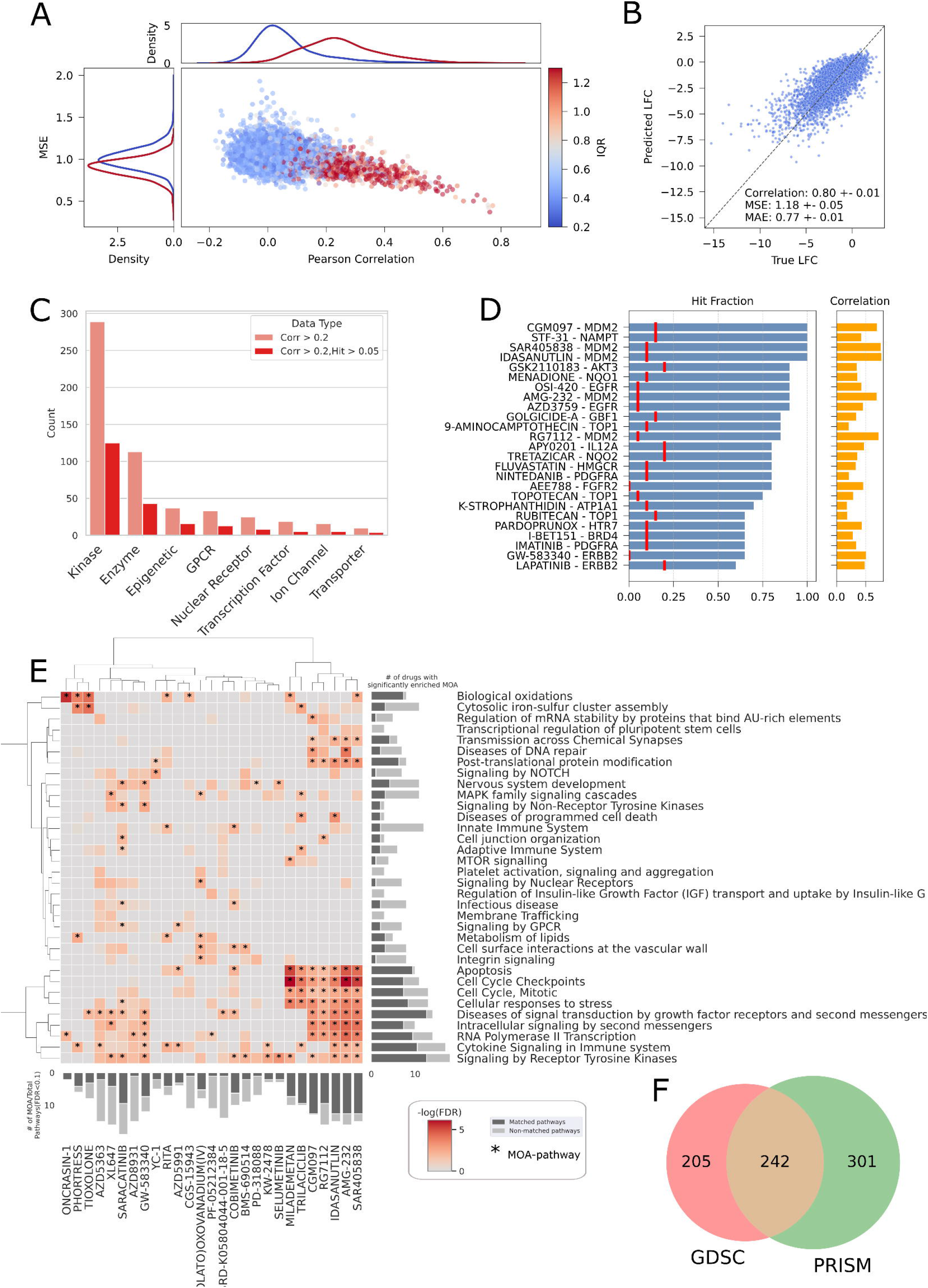
PRISM model performance and interpretation. A) scatter plot of PRISM drug-specific model correlations and MSEs. Dots are colored according to IQR values, ranging from blue to red for increasing values of IQR. The density plots located at the top and left of figure display the distribution of correlation and MSE values for all models in blue and for those models with an IQR greater than 1 in red; B) scatter plot of predicted vs experimental IC50 values from models with correlation > 0.2; C) barplot statistics of PRISM models with corr > 0.2 (salmon) and corr > 0.2 and target recovered, stratified by putative target protein families (red); D) fraction of target recovery for the top 25 ligand-target pairs (same plot as 2A but for PRISM). Red lines represent the 95th percentile of the Hit Fraction distribution across all genes for a given drug. The length of the bars on the right shows the median pearson correlation for each drug-specific model; E) heatmap showing significant MOA-pathways (rows) for drug models (columns) with corr > 0.5 (same plot as 3E but for PRISM); F) Venn diagram comparing MOA-pathway significantly enriched on PRISM’s and GDSC’s drug model important genes.

We obtained a total of 762 drug models with a correlation greater than 0.2 (Supplementary Table 7). The aggregated IC50 predictions of these models with experimental values achieved a ρ=0.80 and MSE=1.18 (Figure 4B). Drugs targeting kinases are by far the category with the highest number of models with a correlation greater than 0.2, being also more effective in recovering the corresponding targets among important genes (Figure 4C). Other recurrent drug target categories are generic Enzymes, followed by Epigenetic and GPCR targets (Figure 4C). Kinases also stand out when normalizing the number of models for the total number of drugs considered in PRISM (Supplementary Figure 2A,B), or considering the number of drugs achieving a certain LFC threshold (Supplementary Figure 2C). However, other drug targets such as Epigenetics regulators and Transcription Factors ranked higher when considering normalized counts due to the lower total number of drugs in these classes (Supplementary Figure 2B). The normalized count metric penalized instead popular drug targets, such as GPCRs or Ion Channels (Supplementary Figure 2B,C). Notably, they were characterized by a very small number of drugs with IQR greater than 1 (Supplementary Figure 2A), suggesting very specific effects on fewer cell lines.

Inspection of the most important genes revealed that 62% (339 out of 547) of the models for drugs with target information identified at least one corresponding target among the important genes across at least one train/test split (Figure 4D; Supplementary Table 8). Also for the PRISM dataset, we successfully identified 73.7% of drug targets at or above the 90th percentile threshold of the background recovery distribution (Figure 4D). STF-31 and CGM097 were the two drug models that recovered their corresponding targets (the regulator of intracellular NAD+ pool Nicotinamide phosphoribosyltransferase, NAMPT; and the p53 ubiquitin ligase MDM2, respectively) in 100% of the train/test splits (Figure 4D). Among the most supported models, we also found a few non-oncological drugs, such as PARDOPRUNOX (targeting the Serotonin 5-HT7 receptor HTR7), which is approved as a treatment for Parkinson.

Similarly to the GDSC drug datasets, we curated MOA-pathways for 6305 PRISM drugs (Supplementary Table 9). We compared the pathways more recurrently associated to those drugs with available category annotations, showing that chemotherapeutic agents have distinct pathway signatures with respect to non-oncological as well as targeted drugs (Supplementary Figure 3). We also performed pathway analysis on important genes for each drug model and similarly displayed enrichments for the best-performing instances (i.e. models with ρ>0.5). Certain pathways, e.g. “Cell Cycle, Mitotic”, “Cellular responses to stress”, “Apoptosis”, “Cell Cycle Checkpoints”, “Cytokine Signaling in Immune System” and “Signaling by Receptor Tyrosine Kinases”, were significantly enriched in a larger set of drugs, consistently with their MOAs (Figure 4E, Supplementary Table 10). Notably, these pathways are largely overlapping with recurrently enriched pathways in GDSC drug models (Figure 3D). These pathways characterize a restricted pool of drugs (Figure 4E, right cluster), while a second, larger pool is also characterized by the enrichment of more specific pathways (Figure 4E, left cluster). Intriguingly, while several enriched MOA-pathways also match processes that are recurrently enriched in GDSC oncology drugs, we found a bigger proportion of pathways exclusively enriched only on PRISM drugs, suggesting a broader diversity of mechanisms of actions (Figure 4F, Supplementary Figure 3A).

### MOA-primed models improve drug-sensitivity predictions

We leveraged the rich information of MOA-pathways to select the most informative variables (i.e. gene expression) for a given drug to develop knowledge-driven models (hereinafter referred to as MOA-primed models).

For model priming, we considered MOA-pathways generated via GPT-4 as well as through the freely available Mixtral Instruct model. Models obtained through the latter approach (i.e. Mixtral Instruct) achieved higher performances (Figure 1B,C) and are discussed further below.

On GDSC, MOA-primed models performed overall better than their all-gene counterparts, with median ρ=0.5 (Figure 5A). Overall, aggregated IC50 predictions have a high correlation with experimental values (ρ=0.89) and the lowest MSE (1.52) (Figure 5B).

**Figure 5:**
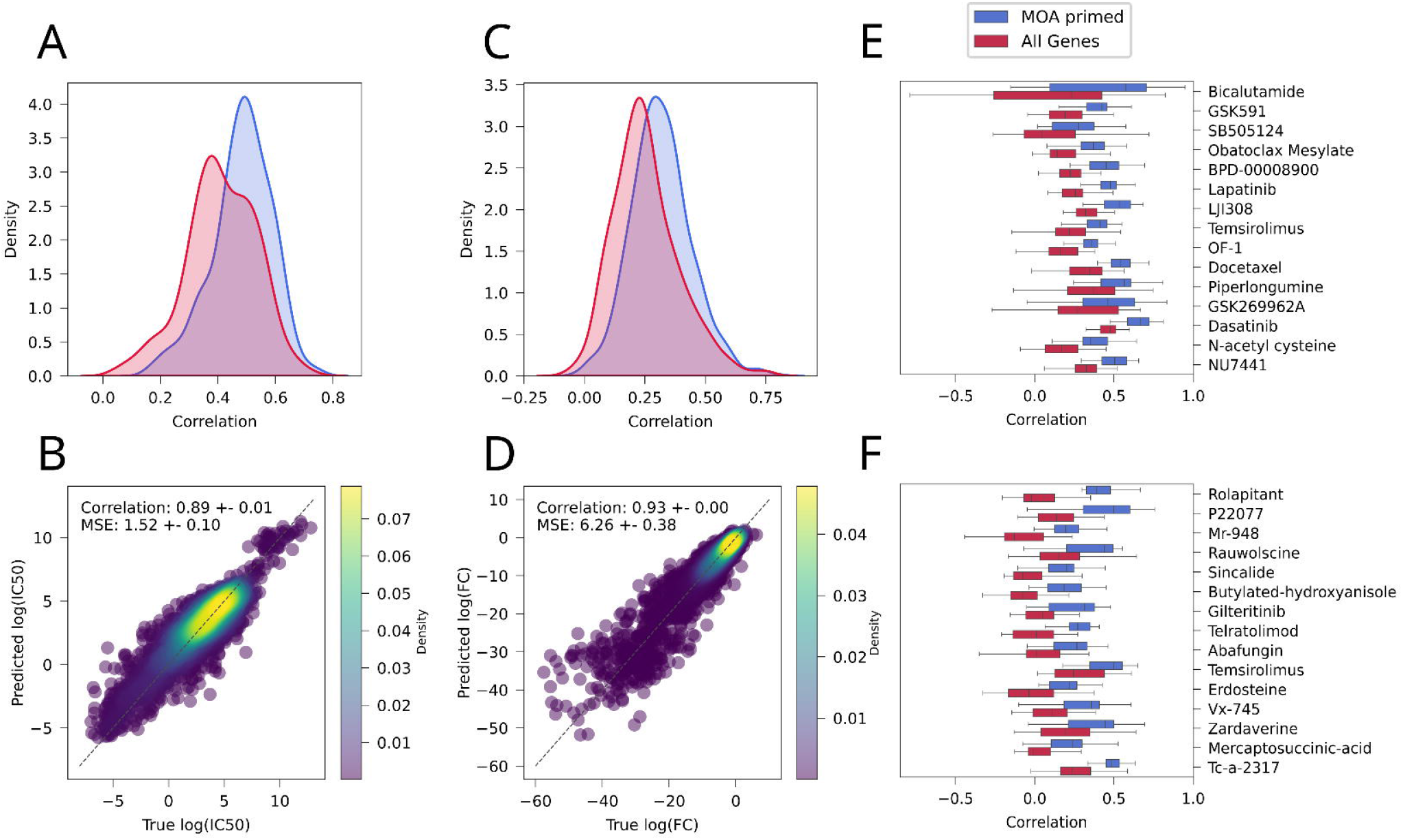
MOA-driven models. A) comparison of the correlation distribution of all-genes (red) vs MOA-primed (blue) GDSC models; B) scatter plot of predicted vs experimental IC50 values from GDSC MOA-primed models. The coloring of the dots on the scatterplot indicates the density of points around a particular area; C) comparison of the correlation distributions of all-genes (red) vs MOA-primed (blue) PRISM models (considering only models with IQR >1); D) scatter plot of predicted vs experimental IC50 values from PRISM MOA-primed models with correlation > 0.2.The coloring of the dots on the scatterplot indicates the density of points around a particular area; E) boxplot of correlation distributions for GDSC models showing the greatest correlation improvement of the MOA-primed vs all-genes models; F) boxplot of correlation distributions for PRISM models showing the greatest correlation improvement of the MOA-primed vs all-genes models

Certain drug models, such as Bicalutamide, showed a remarkable increase in their performance, by almost doubling their correlations when considering only MOA-pathway genes as dependent variables (Figure 5E). On the other hand, a few drug models, such as BX795, Gemcitabine or Savolitinib displayed a reduction in performance of the MOA-primed models compared to all-genes models (Supplementary Figure 4).

Also for PRISM, we employed MOA-pathways to select genes to train MOA-primed models, which outperformed all-genes models. Notably, we obtained almost twice the models with ρ>0.2 compared to all-genes ones (1254 vs 762; Supplementary Table 7). When considering only drugs with IQR > 1, we obtained a median ρ=0.32 (Figure 5C) and a correlation of aggregated IC50 predictions with experimental values ρ=0.96 (Figure 5D). Also in this case, several drugs (e.g. Rolapitant) displayed great improvement in the performance of the MOA-primed model with respect to the all-genes counterpart (Figure 5F).

### CellHit inference on aligned bulk RNAseq from TCGA patients data recovers cancer type-specific mono and combination therapies

We deployed our model to forecast effective drug treatments for TCGA tumors based on their transcriptomic profiles. We first compiled a list of FDA drugs approved for specific cancer types (i.e. https://www.cancer.gov/about-cancer/treatment/drugs/cancer-type) and matched them to the corresponding TCGA cancer types (Supplementary Table 11). This yielded a total of 41 GDSC drugs approved for 23 cancer types. For each drug, we ranked the top 1000 predicted clinical samples according to two criteria: either predicted log IC50 or quantile score (Figure 6A; Supplementary Table 12). The latter is a quantitative measure that trades-off between efficacy and selectivity of a drug, i.e. how much a given drug is predicted to be potent for a particular sample relative to all the other samples inferred (see Methods). In general, both metrics work well in prioritizing patients with matching cancer types (Figure 6B). For certain drugs, such as Cytarabine, Venetoclax, 5-azacytidine we achieved excellent recall statistics for the cancer type for which the drug is prescribed (Figure 6B). Overall, we found that 37 out 41 (90%) of GDSC drugs’ models found, among the top 1k ranked patient samples, at least one from a cancer type for which the corresponding drug is approved (Figure 6C). For several drugs, the majority of predicted samples is derived from cancer types for which that drug has been developed. For instance, Fulvestrant in Breast Cancer (BRCA), BCL2 inhibitors and Cyclophosphamide in Breast Cancer or Acute Myeloid Leukemia (LAML), the mutated BRAF inhibitor Dabrafenib and the MEK1/2 inhibitor Trametinib in skin cutaneous melanoma (SKCM) (Figure 6C). The Dabrafenib inference for SKCM patients is exemplary of the robustness of the model. Indeed, while we don’t find BRAF among the important genes, nor any BRAF-associated pathway among the significantly enriched ones, the model predicts the correct drug for these patients based on gene expression signature similarity, which acts as a surrogate of the mutation status. Indeed, BRAF mutations are the most recurrent among the top 1000 patients scored by the Dabrafenib model (Supplementary Figure 5A), confirming that the model can recognize the mutational status from gene expression signatures. While the majority of the best scored samples with BRAF mutations are affected by melanoma, as expected, we also ranked several BRAF mutated samples from other tumors, such as thyroid carcinoma (THCA) or Diffuse Large B-cell Lymphoma (DLBC) (Supplementary Figure 5B), confirming the repurposing potential of Dabrafenib in more rarely BRAF-mutated tumors (Subbiah et al., 2023). We similarly found for most other drug models that among the top 1k scored samples several from cancer types for which that drug is not currently prescribed (Supplementary Figure 6), suggesting additional possible repurposing candidates.

**Figure 6:**
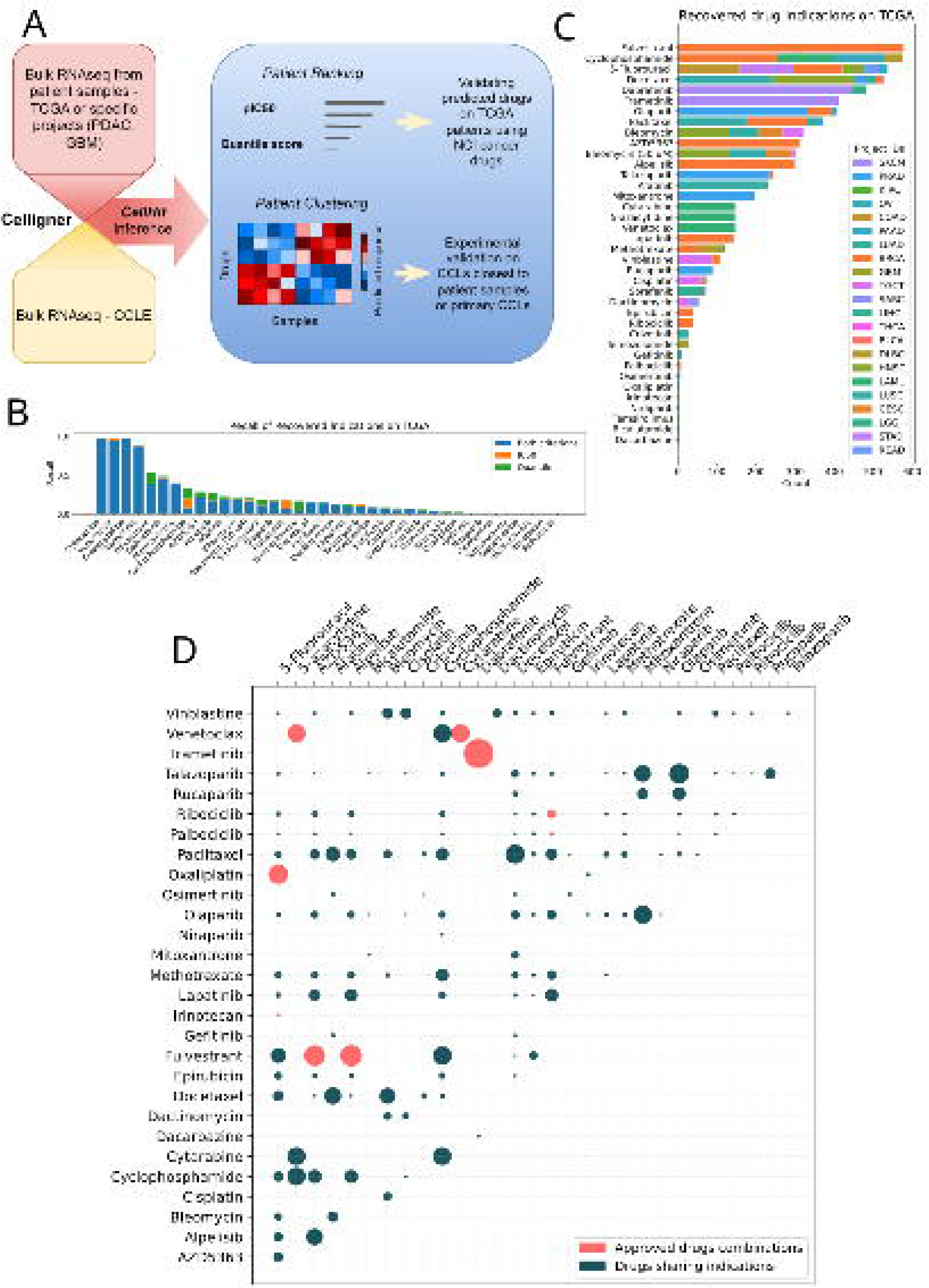
A) Schema of a drug response prediction workflow leveraging Bulk RNA sequencing data from TCGA patients, as well from new PDAC and GBM cohorts. The process begins with data harmonization using Celligner, followed by drug inhibitory concentration (IC50) predictions through CellHit. Patients are then ranked by their predicted predIC50 values and quantile score to assess drug efficacy. Validation involves comparing TCGA predictions with NCI cancer drug metadata and refining tumor-specific predictions by clustering patient responses within cancer subtypes for experimental validation.; B) recall of the recovered drug indications, from NCI cancer drugs, for the TCGA best ranked samples (top 1000), according to either predicted IC50 (predIC50) or quantile score metrics; C) stacked barplot of the GDSC drugs scoring among the top 1000 samples patients with cancer types matching the prescription according to NCI drugs. The height of the barplot’s stacks corresponds to the number of unique samples and the color of the specific cancer type; D) circle plot showing drugs predicted for the same pool of patients, i.e. suggesting combination therapies. Circle diameter is proportional to the number of unique samples, among the top 1k, best scoring for both drug models. Colors indicate the level of support for that combination, i.e., approved (red) or sharing indication for the same cancer type (dark green); E) Inference on TCGA data for the 20 best performing non-oncological drug models in the PRISM dataset. The height of the barplot’s stacks corresponds to the number of unique samples and the color of the specific cancer type (highlighting potential drug repurposing opportunities). Colors are shared with panel C.

A total of 10.5k samples across 33 different cancer types were ranked among the top 1k scoring ones by multiple drug models, suggesting the potential for combination therapies (Figure 6D; Supplementary Table 13). We inspected the predicted drug combinations and ranked them according to the number of samples predicted. This analysis showed that many of the top-ranking combinations are already approved, such as Trametinib and Dabrafenib in SKCM, Venetoclax and Cytarabine or 5-azacytidine in LAML, Fulvestrant and AZD5363, Alpesilib, Ribociclib or Palbociclib in BRCA, as well as Oxaliplatin and 5-Fluorouracil in colon adenocarcinoma (COAD) (Figure 6D; Supplementary Table 14). We found additional predicted combinations characterized by shared indications, highlighting the potential for novel combination therapy (Figure 6D; Supplementary Table 14).

Additionally, we used the models trained on the PRISM dataset to infer drug sensitivities for each TCGA sample by considering the top 1000 samples ranked by each drug model. We considered only the top 20 best performing models for non-oncological drugs, including 8 for Enzymes and 6 for GPCRs (Figure 6E). The analysis revealed specific cancer-type patterns for the best scoring samples for each drug. For instance, two Adenosine Receptors antagonists, i.e. CGS-15943 and MRS-1220, which have been recently proposed as effective therapies against multiple cancers (Arora et al., 2023; Corsello, Spangler, et al., 2020), are indeed predicted for multiple samples of breast (BRCA), liver (LIHC), prostate (PRAD) as well as gastric (STAD) cancers (Figure 6E).

Therefore, we demonstrated that the CellHit models can reliably predict anti-cancer therapies based on patients’ transcriptomics signatures.

### CellHit predictions identified drugs selective for distinct PDAC subtypes

Next, we determined whether CellHit could infer possible drugs with specific effects against selected pancreatic ductal adenocarcinoma (PDAC) subtypes recently identified by a laser microdissection-based spatial transcriptomics approach (Di Chiaro et al., n.d.).

We first verified the projection of the PDAC samples with the cancer cell lines from CCLE database. As expected, PDAC samples were mapped to the tissue of origin at the transcriptome level (Supplementary Figure 7) while they showed different profiles in terms of cancer cell lines responsiveness to drugs (Figure 7A). The Glandular (GL) subtype was mainly associated with the esophagogastric adenocarcinoma cell lines and only in a minor fraction to the pancreatic adenocarcinoma ones. Contrarily, the Transitional (TR) subtype, which is characterized by gene expression programs suggestive of epithelial-mesenchymal transition, was mainly associated with invasive breast carcinoma and head and neck squamous carcinoma. This result suggests that PDAC cancer cell lines may have such heterogenous responses to drugs (Collisson et al., 2011) that the closest cancer cell lines are rarely associated with the pancreatic lineage. The Undifferentiated (UN) subtype(Di Chiaro et al., n.d.), which lacks endodermal gene expression, was excluded for this aim due to the lack of an experimental model useful to recapitulate this PDAC subtype (data not shown).

**Figure 7:**
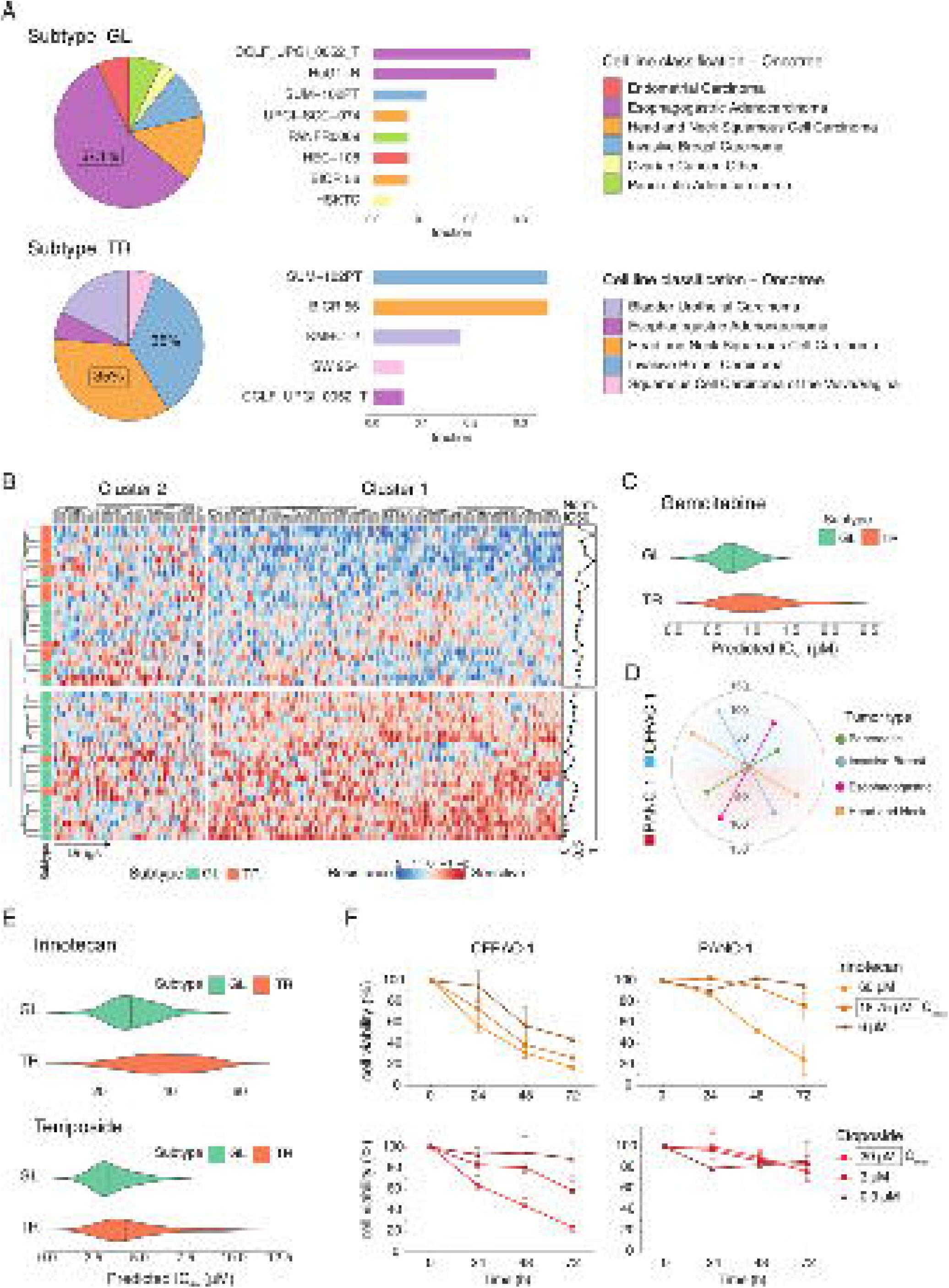
CellHit predictions on distinct PDAC subtypes and experimental validation. A) Enrichment of predicted cancer cell lines that react most similarly to the available drugs for the Glandular (GL) and Transitional (TR) subtypes. Classification of the tissue types of cell lines (oncotree system) is shown; B) Heatmap of predicted IC50 (predIC50) of GDSC drugs derived by CellHit prediction in PDAC samples. K-means clustering (n_clusters=2, method=Euclidean distance) was performed. Subtype annotations are shown for each sample; C) Violin plot showing the predicted IC50 (predIC50) of Gemcitabine for the Glandular (GL) and Transitional (TR) subtypes; D) Euclidean distance derived from Celligner between each PDAC cell line (CFPAC-1, PANC-1) and each selected tumor type (Pancreatic, Esophagogastric, Invasive Breast, Head and Neck); E) Violin plot showing the predicted IC50 (predIC50) of Irinotecan and Etoposide for the Glandular (GL) and Transitional (TR) subtypes; F) percentage of cell viability of CFPAC-1 and PANC-1 cells treated with increasing concentrations of Irinotecan or Etoposide at 24, 48 and 72 hours. Individual values represent the average of three independent experiments ± SD.

We next applied the GDSC-trained version of CellHit to the PDAC samples for the prediction of drugs to which PDAC subtypes showed different sensitivities. Hierarchical clustering based on the predicted IC50 of the available drugs showed that PDAC samples segregated into two main groups according to their subtype of origin (Figure 7B; Supplementary Table 15). Moreover, it also revealed two main clusters of drugs to which PDAC samples showed different sensitivities. In cluster 1, TR samples were associated with resistance to the treatments compared to the GL ones. This finding is in line with the behavior and the poor prognosis of PDACs in which the Transitional subtype is the most abundant tumor component (Di Chiaro et al., n.d.). Cluster 2 was instead enriched in drugs with different effects on both subtypes likely due to the co-existence of endodermal gene programs in these two PDAC variants(Di Chiaro et al., n.d.).

Well-known chemotherapeutic drugs, such as Gemcitabine (cluster 1), were previously described to act mainly on classical epithelial (glandular) cells (Collisson et al., 2011). As expected, CellHit predicted that the GL subtype was more sensitive to Gemcitabine, since these cells have epithelial gene expression programs, compared with the TR subtype which displays quasi-mesenchymal phenotype (Figure 7C). This analysis confirms the subtype-specific responses to this drug (Collisson et al., 2011).

To test these drugs for their repurposing potential, we sought the PDAC cell lines that most closely resembled the cancer cell lines shown by CellHit to have similar drug response profiles to those inferred from patients’ samples (Figure 7A). Celligner revealed that CFPAC-1 was transcriptionally closest to both the pancreatic and the esophagogastric adenocarcinomas while PANC-1 was associated to the invasive breast and the head and neck squamous carcinomas (Figure 7D), lineages that were previously associated by CellHit to GL and TR subtypes, respectively. Among the drugs enriched in cluster 1 and predicted to be more effective in the GL subtype compared to the TR one (Figure 7C,E), we also found two topoisomerases inhibitors, such as Irinotecan and Teposide (or its analogous Etoposide), that are approved and used in clinics for cancer therapy (Fujita et al., 2015; Zhang et al., 2021)(Fujita et al., 2015; Zhang et al., 2021). To validate our predictions, we treated CFPAC-1 and PANC-1 cells with these drugs at different drug concentrations to test the cell viability over three days (Figure 7F). Both treatments were preferentially active in the CFPAC-1 cell line with respect to PANC-1 at the dose corresponding to its Cmax, the maximal concentration that can be reached in patients’ blood. Overall, the experimental validation was in concordance with CellHit predictions confirming the robustness of the model to infer drug responses in different tumor subtypes and underlining the capacity to repurpose FDA-approved drugs for one of the most lethal solid tumors for which there are no efficient drugs available.

### Validation on primary cells from GBM patient’s samples the specific response profiles predicted by CellHit

To further assess *CellHit*‘s capabilities in a real case scenario, we employed the model trained on the GDSC dataset to infer the drug sensitivity profiles for 64 samples obtained from patients with Glioblastoma Multiforme (GBM). We employed the inferred drug sensitivity profiles to cluster patients, which identified two main groups displaying different sensitivity profiles to drugs (Supplementary Figure 8). We chose for experimental validation primary cell lines obtained from the samples Gb130 and Gb107, considered as representatives of the two groups (Supplementary Figure 8). We tested two compounds, the Mcl-1-specific inhibitor AZD5991 and the E3 ubiquitin-protein ligase XIAP inhibitor, AZD5582, as we found them to be respectively more and less sensitive relative to the median values across samples. When considering only the predicted lnIC50, we predicted for both samples Gb130 and Gb107 higher sensitivities for AZD5582 than AZD5991 (Figure 8A), which we experimentally confirmed (Figure 8C,D). Indeed, experimental lnIC50 are in line with the predicted ones, particularly for the Gb130 sample. Predictions of the Gb107 sample differ more compared to experimental results, which anyway confirmed that AZD5582 is more effective in inhibiting the growth of the cancer cell line. Such discrepancy might likely be due to highly specific transcriptional program characterizing this sample and its derived primary cell line, which is closest to a Leiomyosarcoma cell line based on its response patterns (Supplementary Table S16).

**Figure 8:**
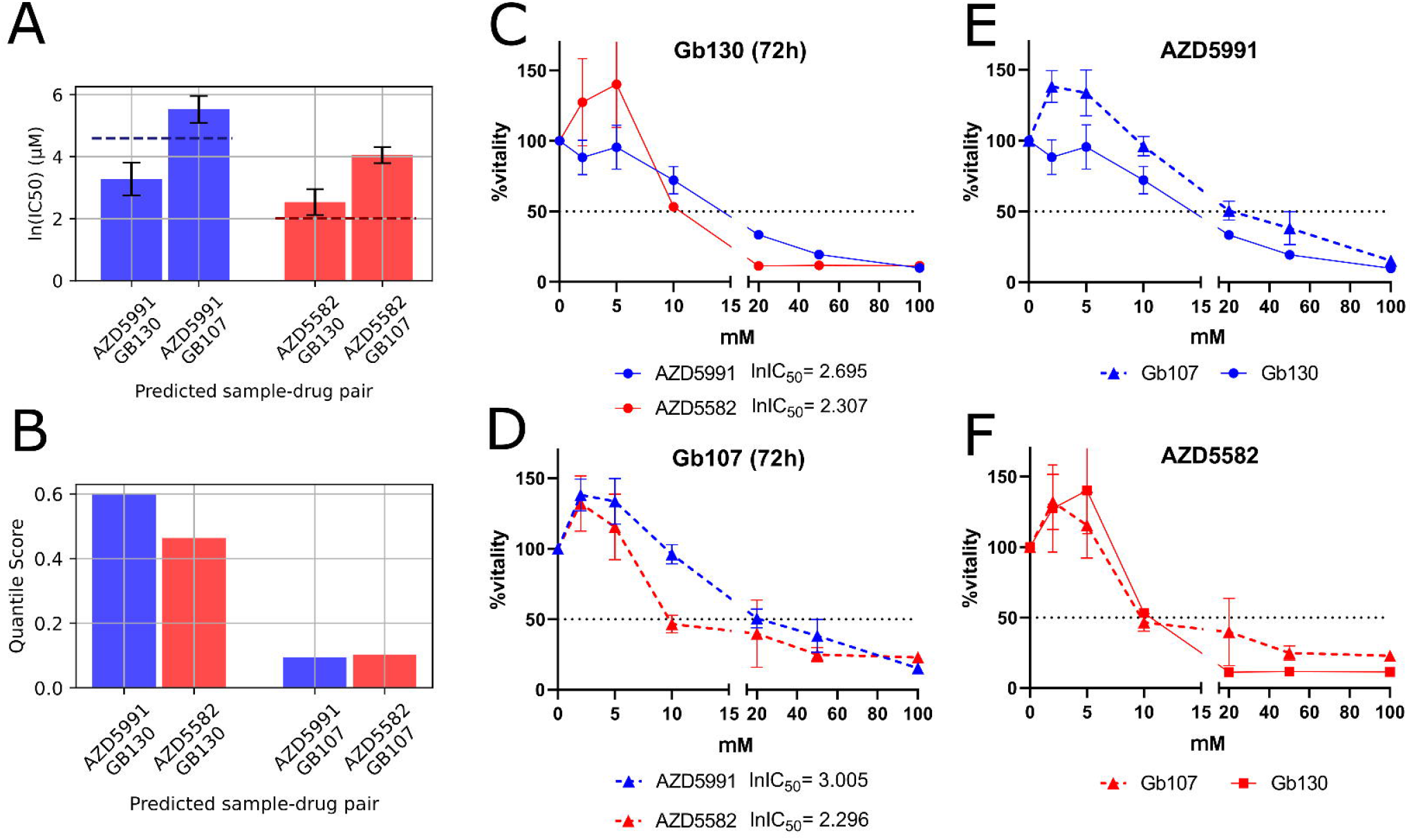
GBM patient samples inference and experimental validation. A) predicted lnIC50 of drugs AZD5991 (blue) and AZD5582 (red) for samples Gb130 and Gb107. Horizontal dashed lines indicate the median of the experimental lnIC50 of each drug on GDSC cell lines; B) predicted quantile scores of drugs AZD5991 (blue) and AZD5582 (red) for samples Gb130 and Gb107; Dose-response curves for the two patient-derived primary cultures of GBM, C) Gb130 and D) Gb107 using the Crystal Violet assay. The curves illustrate the response to treatment in terms of reduction of cell viability with AZD5991 and AZD5582, measured 72 hours post-treatment for Gb107 (C) and Gb130 (D). The IC50 values for both drugs are provided. The dose-response curves are also compared across the two cell lines for AZD5991 (D) and AZD5582 (E). The AZD5991 curve is represented by blue lines, while red lines are used for AZD5582. The profiles for Gb107 are depicted with dashed lines and triangles marking the points, while those for Gb130 are represented with solid lines and circles marking the points. The threshold for a 50% reduction in cell viability is indicated by a dark dashed line.

By considering the deviation of the predicted lnIC50 values from the median of the distribution of experimental lnIC50 across cell lines, it is possible to observe that AZD5991 had a predicted lnIC50 lower than the median lnIC50 (i.e., 4.591) for Gb130 and higher for Gb107, while AZD5582 had a predicted lnIC50 higher than the median lnIC50 (i.e., 2.014) for both Gb130 and Gb107 samples (Figure 8B). Such different extents of deviation from the median are also associated to differences in quantile scores for these drugs for the inferred samples. Indeed, while AZD5991 has a bigger difference of quantile scores among cell lines, with higher values for Gb130, AZD5582 is characterized by more comparable values. We therefore proceeded to test the difference in response of the same drug on the two primary cultures simultaneously (see Methods). We observed a greater reduction in viability in Gb130 compared to Gb107 in response to the treatment with AZD5991 (Figure 8E). This result was consistent with the model, which predicted a lower lnIC50 for GB130 (3.05) than for GB107 (5.57). For AZD5582, we did not observe a clear difference in cell viability reduction following treatment (Figure 8F).

## Discussion

In this study, we have developed a novel ML framework to predict and interpret the sensitivity of cancer cell lines to drug treatments and proposed a strategy to perform robust inference with this model on bulk RNAseq obtained from patients’ samples. Our analysis showed that the critical information to predict cell line drug sensitivity chiefly resides in the features obtained from the cell line and that integration of the drug representation into the model doesn’t significantly improve the predictive performance. This is also expected since the GDSC dataset contains too few chemicals to allow the model to learn any molecular principle associated with their bioactivity. Therefore, we developed individual models for each drug by considering as features only the cell line expression data and showed that their aggregated performance approximates the one of the joint model which also uses drug representations. We found out that XGBoost achieved the best performance compared to other SOTA approaches, such as KRR and SRMF (Chen & Zhang, 2021), or a fully connected MLP neural network.

Importantly, our XGBoost-based model for individual drugs provides much easier means to interpret the predictions and explain which genetic expression features are leveraged by the model. As highlighted by (Corsello et al., 2020), numerous drugs display selective activity profiles. Consequently, a detailed assessment of model performance must include the evaluation of their predictive power in identifying those specific deviations from a drug’s usual cytotoxic or inhibitory effects (i.e., deviations from a drug’s specific mean/median activity). This approach enabled us to move past the biases inherent in the data, providing a more specific depiction of the models’ predictive capabilities. Moreover, the ability to predict drug activity specificities across cell lines has significant biological implications, beyond the statistical advantages already mentioned. Variations in the efficacy of drugs, particularly concerning specific types of cancer, reveal key molecular mechanisms that underpin and distinguish different pathologies. Insights gained from the interpretability of our models shed light on these nuances, enabling a deeper understanding of drug interactions with genetic expression profiles. This knowledge can be pivotal in the development of targeted therapies within a personalized medicine framework, where understanding the particularities of drug responses is essential for tailoring treatments to individual patient profiles. The dual criterion that we have employed for determining feature importance, i.e. SHAP and permutation importance, sets a very stringent requirement for the identification of important expression signatures. Despite this, we found that many drug models are indeed learning the mechanisms responsible for cell-line drug sensitivity solely based on cell-line basal transcriptomics data.

We employed a strategy based on a freely available LLM, i.e. Mixtral Instruct, to improve the MOA description for each drug and use it to associate semantically closest pathways from a reference knowledge base. Through this resource, we assessed that many of the models are characterized by significantly enriched pathways matching the MOA for the corresponding drug, in addition to frequently identifying the nominal targets. We also tracked many significantly enriched pathways that do not match the known MOA for a specific drug, suggesting that additional, new MOAs might support the effect of certain drugs against cancer cells. Overall, 135 GDSC drug models out of 253 can recover either the target or MOA-related pathways among important genes, suggesting that drug-specific models leverage known biological mechanisms to carry out their prediction task. We also demonstrated that the important genes of the model successfully recapitulated cancer tissue essential genes from a recently published compendium (Pacini et al., 2024). Our models’ interpretation clearly suggests that drug sensitivity emerges from the interplay of drug MOAs and genes essential for cell survival. We speculate that the more efficient the cross-talk between MOAs and essential genes, the more effective a drug is in inhibiting cancer cell growth. Our purely data-driven approach to explain drug sensitivity mechanisms can be considered as a complementation of previous approaches either based on simple correlation of sensitivity and basal gene expression (Rees et al., 2016) or integrated with PPI network and pathway analysis (Fernández-Torras et al., 2019).

The proposed LLM pipeline extracts features in a human-readable and interpretable manner, which is crucial in omics problems where the number of features (genes), greatly exceeds the number of available training points (cell lines or samples). The proposed pipeline represents an example of how to use powerful LLMs by leveraging relevant domain knowledge. With the very fast advancements in the field regarding model efficiency and the lowering cost of computing resources, it will probably be possible in the future to obtain deeper and more granular insights. Moreover, the growing ‘reasoning’ capabilities of future models could further improve the capabilities of the proposed pipeline by leveraging multi-modal contents, such as images and knowledge graphs and similar applications of LLMs are already appearing in the literature (e.g. https://biochatter.org/). The proposed interpretability pipeline represents both a novel framework and a toolkit applicable beyond our current datasets to address the complexities of high-dimensional datasets commonly encountered in the “omics” realm. This pipeline bridges the gap between state-of-the-art Explainable AI (XAI) techniques, such as feature permutation and SHAP values, with traditional gene enrichment and over-representation analysis. This integration serves a dual purpose: firstly, it provides a mechanism to delve into the model’s mechanistic underpinnings, offering opportunities for debugging and enhancing our understanding of its internal processes. Secondly, by providing interpretability insights on a multi-scale level— from local to aggregated explanations—it facilitates the formulation of new, experimentally testable biological hypotheses.

In our study, we first tailored the training and interpretability strategies to model the GDSCv2 dataset, which has also been extensively tackled by a multitude of previous methods (J. Chen & Zhang, 2021; Firoozbakht et al., 2022; Xia et al., 2022). We then applied the same strategy to the PRISM dataset, which consists of a much larger panel of drugs screened against cancer cell lines. Although the PRISM dataset poses specific challenges, such as many drugs showing little or no effect at all on cancer cells, we showed that several models achieved good performances and recovered the information of targets and associated MOAs, which encompass a broader range of biological processes compared to GDSC drugs. To the best of our knowledge, this is the first competitive ML model tackling drug sensitivity predictions on both GDSC and PRISM through an explainable framework. However, it is possible that the kind of regression model that we are employing here might not be best suited for many of the drug candidates that we have observed in PRISM, characterized by highly specific activity profiles (in other words, showing activity on a limited number of cells). Considering the high anti-cancer potential of many non-oncological drugs against certain targets (e.g. GPCR drugs) (Arora et al., 2023; Wu et al., 2023), it will be interesting to explore in the future alternative ML frameworks to model the sensitivity of those drugs for which we obtained low performances due to high specificity and low variability of the response.

To deploy the models in real world scenarios, i.e. on bulk RNAseq data obtained from patients, we employed Cellligner(Warren et al., 2021) to align bulk RNAseq from patients to those from cell lines upon which the model has been trained. We used Celligner’s transformed RNAseq data both to train the model (i.e. CCLE) as well as to perform inference on more than 10k patients’ samples from TCGA to predict an IC50 for each drug. Best predicted drugs for each patient often matched mono- and combination-therapies approved for the corresponding cancer type, such as Venetoclax AML, Fulvestrant in BRCA and Dabrafenib in SKCM, along with multiple combination drugs whose usage has been already approved. These results support the high potential for translation of our model predictions, as we find many more predicted mono- or combination-therapies for many TCGA cancer types, with some evidence of indications for combined usage, which might represent new repositioning opportunities.

To further validate our strategy, we transformed RNAseq data from different morpho-biotypes recently identified for PDAC(Di Chiaro et al., 2024) and inferred the most likely drugs for the samples of distinct subtypes. We showed that Irinotecan and Etoposide, predicted to be more effective against the “GL” than “TR” biotypes, indeed displayed differential sensitivity on cell lines more closely resembling the different tumor types reacting most similarly to the two PDAC subtypes, corroborating our predictions. The high intratumor heterogeneity, namely the coexistence in the same patient of heterogeneous groups of tumor cells with distinct morphological and transcriptional profiles, may lead to the adaptation/selection of these cells to therapy. Thus, this approach could be important to design ad hoc combinatorial therapies based on the tumor’s subtype composition. We also demonstrated on GBM patients’ tumor samples the capability of our model to exploit predicted drug sensitivity profiles to cluster samples based on their similarities in drug sensitivity profiles. Through this approach we have identified representative samples with specific sensitivities, which we experimentally validated using match patient-derived tumor primary cell lines. These additional validations not only strengthen the reliability of our model but also highlights its potential translatability in clinical practice providing the therapeutic field of glioblastoma, which has remained stagnant for years, with a broad spectrum of potential new therapeutic possibilities.

These results pave the way for future exploitation of the CellHit model for inference using larger sets of drugs (i.e. PRISM) on alternative patient cohorts. In this respect, it will be critical to develop faster algorithms to align bulk RNAseq for inference with the model. The current strategy, based on Celligner, requires an initial alignment between CCLE, TCGA and any additional input RNAseq dataset, and subsequent retraining of the model on the transformed CCLE data. We plan to employ deep learning architectures, such as Variational Auto Encoders (e.g. Mober(Dimitrieva et al., 2022)), to improve this preliminary alignment step which is of critical importance to effectively deploy cell line-based models on patient samples. We plan to provide our models via a webapp for fast drug-sensitivity inference of bulk RNAseq data from patient samples provided as input. This will allow to analyze and compare inputted samples based on the similarity of their *responsiveness* profile, in addition to the one based on transcriptomic profile, which will speed up the hypothesis generation process to find new personalized treatments against cancer.

## Methods

### Datasets

#### Transcriptomics, IC50s and LFCs

We obtained data model’s training from two comprehensive high-throughput studies: the Genomics of Drug Sensitivity in Cancer (GDSC) (Iorio et al., 2016) and the Profiling Relative Inhibition Simultaneously in Mixtures (PRISM) (Corsello et al., 2020). We downloaded the GDSC2 dataset, release version from 24 July 2022, from its official website (https://www.cancerrxgene.org/). This dataset consists of 969 cancer cell lines profiled for their responses to a panel of 286 drugs. Our primary focus within this dataset are the half-maximal inhibitory concentrations (IC50) values, which serve as target values for predictive modeling. We also incorporated the PRISM Repurposing Public 23Q2 dataset from the Dependency Map (DepMap, Broad Institute) portal, considering log-fold change (LFC) in cell viability measurements and consisting of 919 cell lines and 6.415 compounds. We also retrieved drugs metadata available both in GDSC and PRISM regarding drug putative targets and mechanism of action. We obtained RNA sequencing (RNASeq) data from the Cancer Cell Lines Encyclopaedia (CCLE)(Barretina et al., 2012) as available from DepMap. We considered log2-transformed transcripts per million plus one (TPM+1) data for protein-coding genes. We mapped CCLE cell lines data to GDSC by using available COSMIC to DepMap identifier mappings which allowed us to link cell lines to additional resources available on the DepMap portal such as mutations, gene essentiality, and additional metadata. We employed DepMap IDs to sort cell lines into different tissues and disease categories via the OncoTree classification system(Kundra et al., 2021).

#### TCGA data

We integrated into our pipeline RNA bulk transcriptomic data from The Cancer Genome Atlas (TCGA) (Chang et al., 2013) which we used for initial validation of the designed models on actual patient-derived samples. Specifically, our study employs the Tumor Compendium v11 Public PolyA dataset, released in April 2020, which amalgamates data from various publicly accessible repositories, including the TCGA and Therapeutically Applicable Research to Generate Effective Treatments (TARGET) projects. This data, sourced from the UCSC Treehouse Public Data platform, is downloaded in the RSEM log2(TPM + 1) normalized format.

#### Gene essentiality data

We retrieved data from the 23Q4 CRISPR Gene Dependency dataset from the DepMap portal. This dataset contains synthetic lethality experimental data across 1,100 unique cell lines and 18,444 genes. We refined our analysis to a subset of 17,425 genes, which are also represented in both the CCLE and TCGA databases. Matching and integration of this dataset with cell lines from GDSC and PRISM is made through DepMapID.

#### Mutations data

We obtained somatic mutations data from the 23Q4 release of the DepMap portal. This dataset includes mutation data for 1,750 unique cell lines. We considered the mutations of 693 high-consensus oncogenes and tumor suppressor genes as curated in the OncoKB database(Chakravarty et al., 2017). We binarized oncodriver genes as mutated or not mutated regardless of the specific mutation type. We cross-referenced this dataset with cell lines from GDSC and PRISM via DepMapID.

#### Reactome

Pathway data from the Reactome database(Jassal et al., 2020) is incorporated into our analysis. Notably, we leveraged the directed acyclic graph representing the hierarchical structure of the pathways alongside a comprehensive list of pathways and their associated genes. Data extraction from Reactome is executed through two main methods: file dumps from Reactome, which provide us with the list of pathways and their hierarchical organization, and the REST API, which is used to obtain information on pathway-associated genes and the drugs that have been manually annotated to these pathways. The API is queried programmatically using the “get” function from Python’s built-in “requests” library. We employed topological sorting to organize the nodes (i.e., pathways) of the Reactome hierarchy, considered as a directed acyclic graph (DAG). This sorting technique arranges the nodes in a linear order where each node U, having a directed edge towards node V, precedes V, ensuring that all directed edges progress from higher to lower layers. Through this process, we systematically categorized nodes into hierarchical “layers”, designed to only have incoming edges from the layer above and outgoing edges to the layer below. We focused on pathways within the “1st Layer,” selecting a subset of 169 unique pathways from Reactome. Processing of this data is carried out by leveraging Python’s networkx library, specifically its “topological_sort” function.

#### NCI Cancer drugs

We developed a natural language processing pipeline to programmatically identify drugs’ clinical indications for the different types of cancer as defined in TCGA. In more depth, this pipeline works by extracting information from textual data. Utilizing the Beautiful Soup (https://www.crummy.com/software/BeautifulSoup/bs4/doc/) package, we retrieved links to drugs listed on the National Cancer Institute’s website (https://www.cancer.gov/about-cancer/treatment/drugs). We then combined Python’s requests package with Beautiful Soup to extract textual descriptions of each drug. The extracted text is used as input to create a custom textual prompt. This prompt is then fed to an LLM with generation constrained by the Guidance package (see “Mixtral pipeline” section). The goal was to determine whether the text provided any evidence that the drug was prescribed for any of the 37 cancer categories from TCGA. Given the potential discrepancies in drug naming conventions between NCI and GDSC, we recover PubChem IDs from free-text drug names using PubChemPy, a tool developed by the curators of PubChem(Kim et al., 2016) to programmatically retrieve information about chemical compounds. This step was vital in aligning these drugs with their counterparts in the GDSC, ultimately allowing us to obtain clinical indications for a total of 41 drugs. All data obtained through this pipeline is reported in Supplementary Table S11.

#### PRISM drugs’ metadata and IQR analysis

We obtained PRISM drugs categories, i.e. Chemotherapeutics, Targeted Oncology Drugs, and Non-oncology Drugs, from the secondary screening metadata provided by the 2019 PRISM project data release, available through the DepMap portal (https://depmap.org/repurposing/). We transferred the classification to the drugs in the 2023 release of PRISM by matching their BROAD IDs. This yielded a final set of 835 drugs, including 435 targeted therapies, 346 general oncology treatments, and 54 chemotherapeutics. We calculated the Interquartile Range (IQR) for the Log Fold Change (LFC) data across all 6,337 drugs represented in the PRISM dataset. It is important to note that, due to the properties of the logarithm in base 2, an IQR greater than 1 implies that the viability counts of the cell line at the 75th percentile after a given drug are at least twice as high as those observed for the cell line at the 25th percentile.

#### PDAC subtype data

For the drug prediction in PDAC subtypes (Glandular, GL; Transitional, TR; Undifferentiated, UN), data were obtained from the transcriptional profiles of multiple morphological distinguishable tumor areas isolated by laser micro-dissection (LMD) in primary PDACs of treatment-naïve patients (Di Chiaro et al., 2022).

### Data pre-processing

#### Standardization and filtering

We first selected a subset of common genes among the datasets, using the Human Gene Nomenclature Committee (HGNC) (https://www.genenames.org/) system. We removed genes exhibiting zero standard deviation in either of the datasets (CCLE or TCGA), yielding a final set of 18.174 genes. We standardized transcriptomic data by subtracting the mean and dividing by the standard deviation across the whole dataset. This normalization is not particularly important for tree-based models but is important for the stability of neural networks and other models (used as baselines in this work). For the response values (Ys), we conducted a drug-by-drug standardization, by removing for each drug the mean and standard deviation computed specifically for that drug. This standardization ensured that the model’s predictions were not skewed by the inherent differences in the drug response scales but were instead sensitive to the nuances in response patterns specific to each drug, considering that different drugs can have varying ranges and distributions of response values. The standardization was performed only on the training set. The splitting strategy used throughout the analysis is described in the “Model testing” section. At the end of the preprocessing step, the GDSC dataset comprised 686 unique cell lines, 286 unique drugs and a total of 169.208 drug-cell line pairs (IC50 values). The PRISM dataset comprised 887 unique cell lines, 6337 unique drugs and a total of 3.810.028 drug-cell line pairs (LFC values).

#### Alignment of patients’ bulk RNAseq with Celligner

To harmonize representations of bulk RNAseq transcriptomic profiles from cancer cell lines and patient-derived samples, we adopted the pipeline proposed by the Celligner method(Warren et al., 2021). We employed the data from the Cancer Cell Line Encyclopedia (CCLE) and the Tumor Compendium v11 Public PolyA (see “Datasets” section). Unlike the Celligner publication, in our study we also integrated, together with TCGA, bulk RNASeq data obtained from two additional patient cohorts with pancreatic ductal adenocarcinoma (PDAC) and glioblastoma multiforme (GBM)(see below). This integration is achieved by subsetting all data based on a common set of available genes, concatenating experimental samples with TCGA data, and then executing the cell alignment. We used the Python version of Celligner, available through its GitHub repository (https://github.com/broadinstitute/celligner/tree/master).

### Drugs and cells featurization

We featurized the drugs using their two-dimensional structures. Since the GDSC dataset does not provide the structural identifiers for the tested drugs, we used PubChemPy to retrieve the SMILES representations for each tested compound, for a total of 229 unique drugs. We employed the Extended-Connectivity Fingerprints (ECFP)(Rogers & Hahn, 2010) and ChemBerta fingerprints(Ahmad et al., 2022) for drug featurization. Both ECFP and ChemBerta fingerprints were computed using the MolFeat (https://molfeat.datamol.io/) library. As a control, we also introduced a OneHot embedding technique. This approach produced a 229-dimensional vector for each drug, with the ith position marked as ‘1’ for the *i*th drug and ‘0’ in all other positions. In parallel, we also explored alternative featurization strategies for the cell lines. On one hand, we considered the log2+1 normalized expression values of a total of 18.174 genes. We also explored a dimensionality reduction technique, specifically Principal Component Analysis (PCA), retaining eigenvectors describing 90% of the total variance (395 components). PCA analysis is performed through the “PCA” function from Scikit-learn library ((Pedregosa et al., 2011) and fitted on the log2+1 normalized expression values.

### MOA-pathway annotation

To identify pathways potentially involved in the mechanism of action (MOA) of a certain drug, we implemented a pipeline with three different attribution criteria: an LLM attribution criterion, a pathway membership criterion and a manual Reactome pathway annotation criterion. Below we provide detailed explanations for each of these three criteria.

#### Extending drug MOA with LLM

We employed two distinct Large Language Models (LLMs) for the intelligent extraction and interpretation of drug-related information: GPT-4, a proprietary model developed by OpenAI (https://openai.com/chatgpt), and Mistral Instruct, which is freely available and developed by MistralAI (https://mistral.ai/). Information extracted from the LLMs was leveraged to identify biological pathways likely influencing drug efficacy, which can then be used to select on which genes a model should be trained. The full list of curated pathways for each criterion is reported in Supplementary Table S3 and S9 for GDSC and PRISM respectively.

##### GPT-4 based pipeline

The GPT-4 pipeline begins with the extraction of GDSC’s drug metadata. These primarily include short textual labels describing the mechanism of action or the putative target of each drug (253 drugs out of the 286 available in GDSC). Utilizing a specialized prompt, we engaged GPT-4 to expand this basic metadata into a comprehensive textual description, which elucidates the drug’s mechanism of action and its metabolic pathways in greater detail. We then used the expanded drug description as a second specialized prompt to task GPT-4 with the identification of the top 15 biological pathways likely to modulate the drug’s efficacy as well as to provide a reasoned explanation for the selection of each pathway. The model’s ability to elucidate its reasoning offers valuable insights into the complex interplay between drugs and biological systems, significantly augmenting our understanding of drug responses and offering an entry point to possibly debug the pipeline. Both of these steps are implemented using the OpenAI API, with a specific emphasis on the “function calling” feature introduced in July 2023. This feature is pivotal as it enables us to receive responses in a structured JSON format, adhering to a pre-specified schema and allowing integration in the data pipeline. The selection of pathways by GPT-4 is constrained and informed by the Reactome knowledge base, specifically it is forced to select pathways only among “Level 1” pathways (see “Reactome pre-processing”). This hierarchical approach not only provided a comprehensive overview of biological interactions at various levels of complexity but also allowed us to effectively control the scope of pathways among which GPT-4 operates, balancing the complexity and operative cost of our pipeline.

##### Mixtral pipeline

We introduce an AI curation strategy employing Mixtral Instruct(Jiang et al., 2024), a freely available instruction-tuned 8×7 billion parameter mixture of experts LLM. Notably, the usage of a freely available architecture allowed us to devise a cost-effective and reproducible methodology, addressing computational and accessibility challenges. Our approach guarantees predictive accuracy on par with GPT-4 (or even exceeding) by leveraging three core strategies: structured Chain-of-Thought prompts(Wei et al., 2022) for detailed reasoning, Self-Consistency(Wang et al., 2022) procedures for the minimization false positives, and Retrieval Augmented Generation (RAG)(Lewis et al., 2020) for the integration of validated scientific literature.

Operatively the pipeline unfolds across three phases: initial drug description generation based on metadata, refinement of these descriptions with RAG leveraging PubMed abstracts, and the selection of biological pathways through a self-consistency approach. Initially, drug descriptions are generated using detailed prompts that elicit step-by-step reasoning from Mixtral Instruct, mirroring the GPT-4 pipeline but with enhanced specificity. To further diminish the likelihood of model hallucinations, descriptions are refined by integrating a RAG strategy through the Entrez module of the biopython python library(Cock et al., 2009). This involves systematically retrieving and synthesizing information from the top 10 PubMed abstracts (sorted by relevance) related to each drug. Subsequently, the LLM is tasked with refining these initial descriptions, specifically prioritizing information from the abstracts in the event of conflicting statements and seamlessly integrating any relevant, missing knowledge. In the pathway selection phase, we introduce a self-consistency methodology where Mixtral Instruct is tasked multiple times (with different random seeds) to identify the most relevant biological pathways influencing drug efficacy. By considering pathways identified at least twice across multiple iterations, we significantly diminished the risk of false positives. Due to the absence of function calling capabilities in Mixtral Instruct, we interleave “generate” and “choice” functions from the Guidance library (https://github.com/guidance-ai/guidance) to impose constraints on the LLM’s generative process and obtain structured outputs from the LLM. The use of Guidance not only allows for a controlled selection among predefined pathways but also ensures that the rationale behind each choice is generated before the pathway itself. This enhances the causal coherence of the model’s output and leverages its autoregressive nature.

Operationally, we deploy a GPTQ 4-bit quantized version of Mixtral Instruct, sourced from the Huggingface model Hub (https://huggingface.co/TheBloke/Mistral-7B-Instruct-v0.1-GPTQ). This quantized model configuration strikes a balance between model precision and inference speed, significantly reducing computational costs. Crucially, its compatibility with NVIDIA V100 GPU cards, requiring less than 30 GB of VRAM, enabled us to conduct extensive deployment within our High-Performance Computing (HPC) cluster efficiently. Furthermore, the use of the vLLM (Kwon et al., 2023) library facilitates continuous batching and maximizes GPU utilization during inference for drug description generation and refinement. For the task of pathway selection, we integrated the Huggingface transformers library(Wolf et al., 2019) with the Guidance framework to adhere to Guidance requirements. All prompts utilized throughout the Mixtral Instruct pipeline, alongside the project’s code, will be publicly released.

#### Extending drug MOA with target pathways and drug annotations in Reactome

We also linked drugs with potential Reactome pathways based on their putative targets. Utilizing the Reactome API, specifically the “referenceEntities” endpoint, we extracted entities classified under the ReferenceGeneProduct class for each pathway of “Level 1” (see Reactome Preprocessing). This process yielded a comprehensive list of gene products associated with each identified pathway. A pathway was considered relevant to a drug if its putative target was among the pathway’s gene products. We extracted known ligand associations to Reactome’s pathways, by using the Reactome API’s “referenceEntities” endpoint. Specifically, we retrieved for each of the “Level 1” pathways the objects assigned to the ReferenceTherapeutic class. This step provided a nested list of compounds along with their common names and corresponding identifiers from either PubChem or the Guide to Pharmacology for each pathway.

To merge annotations, we re-employed the PubChemPy pipeline introduced in the “Drugs and cells featurization” chapter. For those instances where the Guide to Pharmacology was the initial source, this pipeline converted identifiers to PubChem format. Since we had previously acquired PubChem IDs for molecules within the GDSC dataset, this streamlined the integration process. Through this approach, we successfully associated pathway information with 66 distinct drugs present in the GDSC. We compared retrieved pathways using the three distinct attribution criteria using the Python matplotlib_venn library (https://github.com/konstantint/matplotlib-venn).

##### LLM pathway recovery probability

We assessed the specificity of the LLM in identifying distinct pathways for various drug categories by utilizing metadata from the PRISM 2019 release (refer to “Metadata on PRISM Drug Categorization”). Our examination spanned different drug categories, focusing on the pathways highlighted by the LLM. For each drug category, we calculated the probability of an L1 pathway (see Data pre-processing) to be selected as relevant by dividing the number of times that pathways appear in that drug category by the total number of drugs within that category. To consolidate and visually represent our findings, we compiled the 25 most commonly identified pathways across all categories, presenting them in an annotated heatmap depicted in Supplementary Figure S3.

### Model training

We employed a dual-strategy training methodology for our predictive models, each addressing unique aspects of the drug-cell line interaction landscape. The first strategy involved a joint drug and cell-line featurization approach under the hypothesis that a dual representation of both drug molecules and cells would lead to a more nuanced and accurate prediction of how various cancer cell lines respond to different drugs. As a second strategy, we employed a drug-by-drug modeling approach, where we focused on examining the effects of individual drugs on cancer cell lines, with particular attention to the transcriptional responses of the cells. This method allowed us to delve into the specific mechanisms through which each drug influences cell behavior, thereby offering insights into drug-specific interactions and responses.

#### Validation and hyperparameter optimization

We describe a rigorous approach for validation and hyperparameter optimization essential for extracting maximum performance from deployed models. To obtain a fair comparison between methods, this process is uniformly applied across various models in our study.

The proposed model-based hyperparameter search procedure leverages the Optuna framework (Akiba et al., 2019). Optuna is a cutting-edge tool for automating hyperparameter optimization and, specifically, deploys a Multi-Objective Tree Parzen Estimator (MO TPE)(Ozaki et al., 2020). This estimator simultaneously optimizes two key metrics: correlation and mean squared error, both evaluated on the validation dataset. By optimizing these metrics concurrently, we ensure a balanced approach to model performance, focusing on both absolute predictive accuracy and the strength of the relationship between predicted and observed values. MO TPE leverages a Bayesian-inspired strategy that is initialized with the evaluation of a specified budget of random hyperparameters. These hyperparameters are sampled from a prior distribution (defined by the user) over the space of possible hyperparameter values. This step is crucial for establishing a baseline understanding of the hyperparameter landscape. Upon completion of these random trials, the procedure shifts to a more targeted approach. Utilizing the outcomes of the initial phase, Optuna calculates a posterior distribution of hyperparameters. This shift marks a transition from exploration to exploitation, where the algorithm begins to concentrate on the most promising areas of the hyperparameter space. These areas are identified based on their density of successful outcomes on the validation set, indicating a higher likelihood of optimal model performance. For the models optimized using Optuna in our study, we adhered to a structured budget for evaluations. This included 100 initial random evaluations, serving as a broad exploration of the hyperparameter space. We then conducted an additional 200 evaluations, which were more focused or “greedy.” This approach, totaling 300 trials, is designed to strike a balance between exploring a wide range of possibilities and honing in on the most effective hyperparameters. We ensure transparency and reproducibility of our methodology by providing detailed information about the prior spaces for the tuned hyperparameters of each model.

#### Baselines from the literature

Our investigation includes a benchmarking of various models that have been recognized as state-of-the-art in the literature (J. Chen & Zhang, 2021).

##### Kernel Ridge Regression (KRR)

KRR emerges as a pivotal machine learning technique, offering a sophisticated blend of ridge regression’s regularization capabilities with the kernel trick’s ability to operate in higher-dimensional spaces. Fundamentally, KRR extends linear ridge regression by incorporating a kernel function, thus enabling the modeling of non-linear relationships without explicitly transforming data into a high-dimensional space. This approach effectively addresses overfitting through the introduction of a regularization term, which penalizes the magnitude of the coefficients, thereby constraining their values and ensuring model simplicity. KRR stands as a benchmark alternative for feature selection, distinct from the approaches proposed via Large Language Models (LLMs). This method solely utilizes transcriptomic data, constructing a separate model for each drug without incorporating drug featurizations. We implemented KRR through the Scikit-Learn library.

##### Similarity-Regularized Matrix Factorization (SRMF)

The Similarity-Regularized Matrix Factorization (SRMF) method (J. Chen & Zhang, 2021) is an innovative approach for predicting anticancer drug responses in cell lines. It leverages the inherent similarities between drugs and cell lines to enhance prediction accuracy. Specifically, SRMF incorporates chemical structure similarities of drugs and gene expression profile similarities of cell lines as regularization terms in the matrix factorization model. We implemented this model by modifying the original Matlab code available at (https://github.com/linwang1982/SRMF)

#### Full joint models

Following methodologies that are widely recognized in the literature, we developed multiple predictive pipelines intended to effectively extrapolate and harness meaningful information from both the drugs and the cell lines. Crucially, models in this chapter make use of the featurization described in the “Drugs and cells featurization” chapter.

##### Multi-layer perceptron

We implemented a customizable multi-layer perceptron (MLP) model architecture. The model dynamically constructs its architecture based on specified hyperparameters, including the number of input features, the number of hidden layers, the number of neurons in each hidden layer, and the dropout rate to mitigate overfitting. Each hidden layer is normalized using batch normalization to enhance stability and employs the ReLU activation function to introduce non-linearity, with an optional dropout applied based on the specified rate. The MLP is optimized using the AdamW optimizer, leveraging hyperparameters such as learning rate and weight decay for regularization. Training involves a customizable loss criterion, defaulting to Mean Squared Error Loss for regression tasks, with early stopping implemented based on a patience parameter to prevent overfitting by halting training if the validation metric does not improve for a specified number of epochs. Optimal hyperparameters for this model are fixed following the procedure described in the “Validation and hyperparameter optimization” chapter. The full list of hyperparameters and the prior space of the hyperparameters is provided in the supplementary material. The MLP is implemented using the PyTorch library (Paszke et al., 2019).

##### XGBoost

XGBoost (T. Chen & Guestrin, 2016) stands as a premier machine learning algorithm within the domain of ensemble methods, distinguished by its exceptional efficiency, flexibility, and capacity to deliver high-performance models. Central to XGBoost’s methodology is the concept of gradient boosting, wherein it sequentially constructs a series of decision trees, each designed to correct the errors of its predecessors. This iterative refinement process harnesses the power of gradient descent to minimize a specified loss function, effectively enhancing predictive accuracy with each step. XGBoost further elevates the gradient boosting framework by integrating several key innovations: a regularization component that mitigates overfitting by penalizing complex models, an efficient tree pruning strategy that determines the optimal tree size during the growth phase rather than after its construction, and a scalable handling of missing data. Additionally, it incorporates a novel splitting algorithm capable of evaluating the potential of different tree structures rapidly, thereby optimizing computational resources and facilitating the handling of large-scale data sets. Crucially, given the high dimensionality of our dataset, we leverage the GPU acceleration included in XGBoost from version 2.1.0. Optimal hyperparameters for this model are fixed following the procedure described in the “Validation and hyperparameter optimization” chapter. XGBoost is implemented through the official library (https://xgboost.readthedocs.io/en/stable/)

#### Drug-by-drug models

We delineate the methodology employed for developing a model to predict cancer cell line responses to drugs, focusing exclusively on transcriptomic data. The following methodologies were applied to both the GDSC and the PRISM dataset, resulting in the development of 286 models for GDSC and 6,337 models for PRISM. Crucially, we develop two different types of drug-by-drug models: all genes models and MOA-primed models.

##### All genes models

Given the consistently superior performance of the XGBoost algorithm, as evidenced by our results and a multitude of studies in the literature on tabular data, we have selected it as our primary tool for developing drug-specific models. Details about this model type can be seen in the dedicated “XGBoost” chapter.

##### MOA-Primed models

MOA-primed models differ from regular full-gene models in two key aspects: firstly, they are trained exclusively on a subset of genes identified as relevant to the drug’s MOA by the three criteria described in the chapter “MOA pathway annotation.” Reducing model training to this subset of genes leads to a significant reduction in the number of genes from 18.174 to an average of 4.117, achieving a factor of reduction of 4.4. Secondly, this narrower gene focus allows for a more comprehensive optimization process within a reasonable computation time. In more detail, the MOA-primed model consists of an ensemble of three XGBoost models (each comprising five parallel trees) trained on different subsets of the training data. This approach, along with the reduced number of genes used, greatly decreases the overfitting problem both during hyperparameter optimization and training. As for other models, optimal hyperparameters are fixed following the procedure described in the “Validation and hyperparameter optimization” chapter. Differently from other models, we leverage a 3-fold cross-validation strategy during hyperparameter search. Crucially, each XGBoost model belonging to the ensemble is trained on a different train-validation split.

### Model testing

#### Data splitting procedure

To guarantee a robust evaluation of the predictive performance of our models downstream, we have implemented a strategic approach to partitioning the dataset. This partitioning is guided by the OncoTree classification system, which offers a detailed categorization of cell lines across different tissues involved in cancer. We selected two distinct cell lines from each tissue type specified in the OncoTree classification. This diverse selection ensures a representative cross-section of cancer types, aiding in the generalizability of the model. Subsequently, all drug-cell line pairs related to these selected cell lines were systematically assigned to either the training, validation, or test sets. Such an approach ensures that each set reflects a broad spectrum of tissue types and their corresponding responses. To ensure a reliable estimate of our trained models’ performance, we repeated the train-validation-test procedure described earlier 20 times, each with a different random seed. We use random seeds from 0 to 19. The median values of the metrics obtained, along with their standard deviations, are reported.

#### Aggregated evaluation

We conducted a comprehensive comparison of all trained models by calculating the Mean Squared Error (MSE) and Pearson correlation coefficient for the predicted values across all drug-cell line pairs (encompassing all drugs and cell lines) in both the GDSC and PRISM datasets. It is important to note that baselines, full joint models, and drug-by-drug models adhere to the same standardization schema outlined in the section “Data Pre-processing.” As such, before evaluating the metrics on the test set, we adjusted both the experimental and predicted values (Ys) to their original scales on a drug-by-drug basis. The cumulative predictions of the drug-by-drug models resulted from pooling forecasts made by individual models.

#### Drug specific evaluation

We assessed the models’ local performances by computing the median and standard deviation of both MSE and Pearson correlation coefficient across 20 random splits for each drug-specific model. This evaluation is conducted without reverting the standardized data to its original scale. Crucially, this approach assessed a model’s predictive power around its mean activity value (either IC50 or LFC). Additionally, we derived a final summary metric by calculating the median and standard deviation of all these single-model performances, providing a concise overview of model efficacy in a drug-centric context.

### Model Interpretability

#### Importance computations

We have adopted a stringent criterion to identify genes that are pivotal in our models’ predictions of cancer cell line responses to drugs. To determine the importance of genes, we utilized two distinct but complementary methods(Molnar et al., 2020): SHAP (SHapley Additive exPlanations)(Lundberg & Lee, 2017) importance and Permutation Importance(Breiman, 2001). We imposed that a gene is relevant if both methods yielded importance values greater than zero. Notably, SHAP values provide insights into how each gene contributes to the model’s final prediction. This method effectively quantifies the impact of each variable (gene) in the context of the model’s decision-making process. On the other hand, Permutation Importance evaluates the influence of each gene on the overall performance of the model. This is achieved by measuring the degradation in model performance when the values of a gene are randomly shuffled, thereby disrupting the gene’s original relationship with the response variable. In the results discussed above we chose as reference metric for permutation importance a drop in correlation (we provide in the released code also computations for MSE). TreeSHAP, as implemented in the Python SHAP library, was employed to ascertain the importance of each gene (https://shap.readthedocs.io/). Notably, this method offers a fast and exact computation of otherwise costly SHAP importances for tree-based models. Given the high dimensionality of our dataset, encompassing almost 19k genes, we designed a computational pipeline to expedite the computation of Permutation Importance. Initially, we employed XGBoost built-in feature importance methods based on impurity decrease, a standard computation in tree-based methods, to significantly narrow down the list of potential genes of interest. This approach allowed us to reduce the number of genes to be evaluated for importance by at least a factor of ten. To further accelerate the process, we batched multiple variable computations by designing a Numba-enhanced routine and then performed joint fast inference leveraging the GPU-accelerated implementation of XGBoost. Feature importance computations were conducted individually for each of the 20 random seeds. While the SHAP mean importance values were computed exactly, the permutation feature importances were derived by randomly shuffling a gene’s values and observing the resultant drop in model performance. A gene’s permutation importance, for a given drug and a fixed seed, is obtained by averaging importance values obtained across three independent repetitions of the shuffling procedure. This methodology allowed us to mitigate aleatoric uncertainties inherent in the shuffling process, ensuring a more reliable identification of genes that are genuinely influential in predicting drug responses in cancer cell lines.

#### Putative and pathway gene recovery

In evaluating the soundness of our models, we investigated their ability to identify known putative targets and associated genes (i.e., those within the same pathway as the putative targets). For each random seed, we selected genes with non-zero importance value (as outlined in the “Importance Computation” section) and then calculated the frequency with which each putative gene appears in this subset of significant genes. Additionally, we conducted a parallel analysis involving genes functionally linked to putative targets through pathways. For each random seed and corresponding putative target, we aggregated all genes from the pathways involving the targets. We then determined the overlap between this aggregated set and the significant genes, documenting both the quantity and specific identities of the genes identified. Similar to the approach for putative target identification, the rigor of the selection criteria can be adjusted by focusing on pathway genes that consistently emerged across various random seeds. We refined our evaluation by calculating the empirical distribution of recovery rates for each drug, starting with the importation and binarization of gene importance scores—SHAP values (converted to 1 if greater than 0) and permutation importances (set to 1 if less than 0). By intersecting genes with both metrics equal to one, we identified those of dual significance. A baseline distribution of recovery frequencies across splits was established by averaging these intersected, binarized importances. This drug-specific procedure allowed us to derive critical quantiles (90th, 95th) as benchmarks for significant recovery. We then compared the putative targets’ recovery rates to these percentiles.

#### Pathway enrichment

We performed pathway enrichment on the list of important genes (see above section) identified by each All-Genes model using the GSEAPY (v 1.0.6) package(Fang et al., 2023). We performed an over-representation analysis using the *enrichr* function, considering as categories the “Reactome_2022” pathways from the MsigDB library(Liberzon et al., 2015), version 2023.1.Hs. We considered all the pathways with FDR < 0.1. To check if an enriched pathway represents a mechanism of action of the corresponding drug, we checked if it is connected to the MOA-pathways that we compiled for each drug by traversing the graph of Reactome pathway hierarchy using the *python*-*igraph* (v 0.9.10) library. If the enriched pathway is either a parent or a child of any MOA-pathway for that drug, it is considered to match the MOA for that drug. Enrichment analysis was performed for both GDSC and PRISM. Pathway enrichments obtained through this analysis are reported in Supplementary Tables S4 and S10 for GDSC and PRISM respectively. Enrichment data obtained is also used to assess the proportion of MOA pathways that are overrepresented across the three criteria (refer to the “MOA Pathway Annotation” section). To generate Figure 3C, we only considered the models exhibiting a correlation exceeding 0.5.

We compared significantly enriched pathways on GDSC and PRISM models using the Python matplotlib_venn library (https://github.com/konstantint/matplotlib-venn).

#### Clustering analysis of drug and MOA-related pathways

To elucidate the intricate relationships between drug MOA and their biological pathways, we clustered pathway enrichments and applied stringent statistical filters to identify and visualize significant drug-pathway associations. We focused on drug models with a correlation greater than 0.5 and the pool of MOA pathways found enriched, with a FDR below 0.1, in at least one drug model (see “Pathway enrichment”). For the clustering approach, we employed hierarchical clustering to organize both the drugs and the MOA pathways based on their patterns of association. This method allows for the grouping of drugs with similar pathway profiles and, conversely, pathways commonly influenced by a cohort of drugs. The clustering was conducted using the Ward method within the *clustermap* function from the seaborn python library (https://seaborn.pydata.org/, v0.12.2). The Ward method minimizes variance within clusters, thereby enhancing the interpretability of complex relationships by showcasing clusters of drugs and pathways with similar biological effects.

#### Correlating IC50s, predicted IC50s, Gene Expressions and SHAP values

We aimed to elucidate the relationship between IC50 values, predicted IC50s, gene expressions, and SHAP values to better understand the dynamics of drug response in cell lines. For a given drug, we sorted the IC50 values from the lowest to the highest. This sorting strategy enabled simultaneous comparison of actual versus predicted IC50 values, gene expression levels, and SHAP values corresponding to the gene expression across cell lines. The result of this analysis is depicted in Figure 2C, specifically focusing on the interaction between the Venetoclax drug and the *BCL2* gene. This figure serves to visually convey the complex interplay and correlations among the variables of interest, providing insights into the predictive model’s predictions and the gene’s influence on drug sensitivity. To improve the graphical representation and clarity of trends within the gene expression and SHAP value data, both datasets were subjected to smoothing using a 5-lag moving window.

#### Gene essentiality analysis

We assessed the potential of local SHAP values, representing gene impacts on drug-cell line pair predictions, to identify genes involved in gene essentiality, as evidenced by CRISPR knock-out screenings. Utilizing a dataset from Pacini et al. (Pacini et al., 2024), which lists essential genes across tissues and CRISPR Gene dependencies (see “Datasets”), we tailored this analysis to include only those cell lines and tissues mentioned therein. This required harmonizing tissue nomenclature differences between our dataset and Pacini et al., notably merging Esophagus and Stomach categories into a single group and mapping other tissues’ names to match our dataset’s structure (the mapping will be released together with the code to reproduce the analysis). Our analysis was further refined to drug-cell line pairs exhibiting the highest sensitivity, based on IC50 values and quantile score (see “Quantile Score”). This results in retaining 266 unique drugs and selecting 43,174 “top pairs.” For these pairs, we examined the top k most negative SHAP values to identify influential genes, with *k* set at 10, 20, 50, and 100 thresholds. We remark on how negative SHAP values indicate genes whose inhibition likely enhances drug efficacy, aligning with the biological outcomes expected from CRISPR knockouts that lead to reduced cell viability. By aggregating these key genes and comparing them against the catalog of essential genes from the reference genes, we aimed to ascertain the extent to which local importance estimates could recover genes critical for cell survival and implicated in gene essentiality. We constructed protein-protein interaction networks using STRING(Szklarczyk et al., 2019) within the Cytoscape platform(Lotia et al., 2013). Nodes within the network were annotated with degree information and a cumulative SHAP value pooled from the set SHAP values over ‘top pairs.’

#### Responsiveness similarity analysis

After GDSC model training, we characterize cell lines through a 286-dimensional vector representation derived from the aggregated predictions of these models. This defines a “response space” which complements transcriptomic similarity analyses and can be used to compare samples based on their drug responsiveness similarity. This approach enables two key applications: clustering of cell lines to identify groups with similar drug sensitivities and the use of the Python library *faiss* for efficient nearest neighbor searches in the response space.

### Model Inference on TCGA

#### Quantile score

IC50 values are a commonly used measure of drug potency. However, raw values may not fully capture the nuances of drug efficacy and specificity, particularly for cytotoxic drugs, which could feature systematically lower IC50 levels. We propose a new scoring system aimed at providing a more comprehensive assessment of a drug’s potential.

Our novel approach integrates two dimensions: drug efficacy and drug specificity. Drug efficacy is defined as the likelihood of identifying an alternative drug within the dataset that possesses a higher IC50 value than the current drug in question. This measure reflects the drug’s ability to inhibit or kill cancer cells at lower concentrations, thereby indicating its potency. On the other hand, drug specificity evaluates the probability of finding another cell line upon which the given drug exhibits a higher IC50 value. This dimension captures the drug’s targeted action, ensuring that it effectively inhibits the growth of specific cancer cells with minimal effects on others, thereby reducing potential side effects.

To synthesize these dimensions into a singular, coherent score, we employed the harmonic mean of the two probabilities. The harmonic mean, a type of average, is particularly suited for our purpose because it tends to skew towards the lower of its inputs, ensuring that both high efficacy and high specificity are necessary for a drug to achieve a high score. As such, our metric ranges between 0 and 1, where a score closer to 1 indicates a drug that is both effective and specific. This property of the harmonic mean serves to penalize drugs that are highly effective against a broad range of cell lines (potentially indicating high toxicity) or highly specific but with limited overall efficacy.

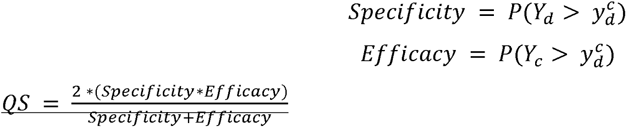

#### TCGA inference and cancer-specific drug prescription validation

We aimed to validate our models using the TCGA dataset. This allowed us to assess whether the proposed pipeline can accurately identify tumor-specific clinical drug prescriptions annotated in the GDSC through the use of processed NCI indications (refer to the “Dataset” section). A key step in our approach was aligning TCGA data with CCLE using the Celligner method (detailed in “Alignment of patients’ bulk RNAseq with Celligner”), which enabled model training on CCLE data and subsequent deployment to TCGA. Moreover, an important part of this validation was to stratify TCGA patients by cancer type, utilizing the extensive metadata available within the TCGA dataset. Our analysis encompassed 9,805 TCGA samples for 41 clinically relevant drugs, with models’ predictions ranked by logIC50 values and quantile scores. We specifically examined the predictive accuracy for the top 1000 patients with the lowest IC50 or highest quantile scores to determine if the highest-scoring predictions correspond to actual clinical prescriptions. This approach effectively measured our models’ clinical prediction reliability. We remarked how TCGA serves as a novel and distinct data source from the training set, representing an unbiased evaluation of our model’s capabilities.

To account for the non-uniform distribution of tumor types across TCGA patients, we augmented our validation with a recall analysis. This additional step aims to determine the fraction of samples associated with each drug’s indication that the models successfully identify, complementing the previous count-based analysis. By stratifying this analysis based on whether samples were identified through their IC50 values, quantile scores, or a combination of both criteria, we ensure a comprehensive evaluation of our models’ ability to accurately predict drug responses. Data from this analysis is reported in Figure 6B. A similar analysis is carried out also on the PRISM dataset. We first ranked the models by performance, then curated a list of the top 20 non-oncological drug models. For each model, predictions were ordered by Log Fold Change (LFC) from lowest to highest, and the initial 1,000 samples were selected. These samples were stratified by tumor type within the TCGA patient dataset to identify potential drug repurposing opportunities.

In conclusion, utilizing the same pipeline, we identified and quantified patients within the TCGA dataset where specific drug pairs are concurrently recommended, indicating potential synergies. We corroborated our findings by cross-referencing manually with the approved drug combinations in DrugBank(Knox et al., 2011), successfully identifying both known and approved combinations.

### Experimental validation on PDAC samples

#### Ranking PDAC cell lines for selected tumor types with Celligner

To determine which PDAC cell lines most closely resemble the tumor types (Pancreatic adenocarcinoma, Esophagogastric adenocarcinoma, Invasive breast carcinoma and head and neck squamous carcinoma) derived from Fig 7A, Celligner (https://depmap.org/portal/celligner/) was used. For each of these tumors, we ranked and selected PDAC cell lines based on their Euclidean distance from that tumor type. The Euclidean distance was calculated as the average distance between a cell line and all TCGA tumor samples of that tumor type.

#### Cell lines, culture and treatments

The following human PDAC cell lines were used: CFPAC1 (KRAS G12V; ATCC CRL-1918) and PANC1 (KRAS G12D; ATCC CRL-1469), were used for the following experiments. CFPAC1 cells were maintained in Iscove’s Modified Dulbecco’s Medium (IMDM) +10% fetal bovine serum (FBS) while PANC1 were maintained in Dulbecco’s Modified Eagle Medium (DMEM) +10% fetal bovine serum (FBS). Media were all supplemented with 2 mM L-glutamine. The cell line was authenticated by the IEO Tissue Culture Facility using the GenePrint10 System (Promega) and was routinely screened for Mycoplasma contamination. Irinotecan (Sigma-Aldrich, I1406) and Etoposide (Sigma-Aldrich, E1383) were diluted in DMSO and used at 3 different concentrate ions for the cell viability experiments.

#### Cell Viability Assays

5-10,000 cells per well were plated in 96-well plates. One day after, cells were treated with the drug (Irinotecan or Etoposide). The cell viability was measured after 72h of treatment using CellTiter-Glo Luminescent Cell Viability Assay (Promega, G9242) and GloMax® (Promega). The assay was performed three times in triplicates.

### Experimental validation on GBM samples

#### Human Subjects

Tumor specimens were obtained from 64 patients who underwent surgical resection of histologically confirmed GBM after providing informed consent. Samples were acquired from the Unit of Neurosurgery of Livorno Civil Hospital. All patients had a GBM diagnosis without a prior history of brain neoplasia and did not exhibit R132 IDH1 or R172 IDH2 mutations. Resected tumors were preserved in MACS tissue storage solution (Miltenyi Biotec, Bergisch Gladbach, Germany) at 4°C for 2–4 hours. Each tumor specimen was rinsed with Dulbecco’s phosphate-buffered saline (DPBS) (Gibco, Carlsbad, CA, USA) within a sterile dish and divided into ∼ 0.5-1 mm2 pieces under a biological hood. Biopsies were cryopreserved in 90% fetal bovine serum (FBS) and 1% dimethyl sulfoxide (DMSO) at −140°C until further preparations.

#### RNA Isolation

RNA extraction from GBM tissues was performed using the Maxwell 16 LEV Simply RNA Tissue Kit (Promega, Madison, WI, USA) according to the manufacturer’s protocol. The concentration of RNA was assessed using the Qubit Fluorometer (Thermo Fisher Scientific, Waltham, MA, USA), and quality was evaluated using the Agilent 2200 Tapestation (Agilent Technologies, Santa Clara, CA) system.

#### Whole Transcriptome RNA Analysis Libraries

RNA-Seq was performed on the NextSeq 500 platform (Illumina, San Diego, CA, USA). Libraries were prepared using the Illumina Stranded Total RNA Prep with Ribo-Zero Plus kit (Illumina), starting from 200 ng of total RNA, following the manufacturer’s protocol. The libraries were quantified using Qubit reagents (Thermo Fisher Scientific, Waltham, MA, USA) and analyzed for validation using TapeStation (Agilent Technologies, Santa Clara, CA). Up to 10 libraries were loaded onto the NextSeq High Output cartridge (150 cycles).

#### Human Subjects for GBM Primary Cell Cultures

Tumor specimens were obtained from 2 patients who underwent surgical resection of histologically confirmed GBM after providing informed consent. Samples were acquired from the Unit of Neurosurgery of Livorno Civil Hospital. All patients had a GBM diagnosis without a prior history of brain neoplasia and did not exhibit R132 IDH1 or R172 IDH2 mutations. The patients included one female and one male, aged 37 and 76 years, respectively. Resected tumors were preserved in MACS tissue storage solution (Miltenyi Biotec, Bergisch Gladbach, Germany) at 4°C for 2–4 hours. Each tumor specimen was rinsed with Dulbecco’s phosphate-buffered saline (DPBS) (Gibco, Carlsbad, CA, USA) within a sterile dish and divided into ∼ 0.5-1 mm2 pieces under a biological hood. Biopsies were cryopreserved in 90% fetal bovine serum (FBS) and 1% dimethyl sulfoxide (DMSO) at −140°C until further preparations.

#### GBM Primary Cell Cultures

Patient-derived GB living tissues were dissociated into single-cell suspensions using mechanical dissociation combined with enzymatic degradation, utilizing the Brain Tumor Dissociation Kit (Myltenyi Biotech, Bergisch Gladbach, Germany), following the manufacturer’s instructions. The culture medium consisted of DMEM/F12 medium (Gibco, Carlsbad, CA, USA), supplemented with 10% FBS (Gibco, Carlsbad, CA, USA), 100 U/mL penicillin, and 0.1 mg/mL streptomycin (Gibco, Carlsbad, CA, USA), 1% Amphotericin B (Gibco, Carlsbad, CA, USA), 1% G-5 supplement (Gibco, Carlsbad, CA, USA), and 1% Glutamax (Gibco, Carlsbad, CA, USA). Cultures were maintained in a 5% CO2 atmosphere at 37°C. The medium was replaced the day after culture. Upon reaching confluency, cells were seeded for viability assays, WST-1, and Crystal Violet, as described below.

#### Treatments of GBM Primary Cell Cultures

AZD5591 (Aurogene S.r.l, Rome, Italy) and AZD5582 (Sigma, St. Louis, MI, USA) were dissolved in DMSO and water, respectively, to create stock concentrations of 200 mM and 4.8 mM, respectively. Dilutions of these drugs for treatment were prepared with the cell medium. We used concentrations of 2, 5, 10, 20, 50, and 100 uM of AZD5991 or AZD5582, or an equivalent volume of vehicle (DMSO or water) for control groups.

#### Viability and Cytotoxicity Assays

We used the Crystal Violet (CV) assay to evaluate cytotoxicity of the same chemicals on the 2 different cell types. For this assay, 2500 cells were seeded in 48-well plates (3 wells per experimental condition). After 72 hours post-treatment, cells were fixed with 4% PFA and stained using a crystal violet solution (0.1% crystal violet, 20% methanol, in water). Following staining, the excess crystal violet was washed with tap water, and plates were dried. Cells were de-stained using a 10% acetic acid solution, and the absorbance of the solution was then measured at 590 nm. IC50 values were computed using python’s SciPy, library reimplementing the procedure of Corsello (2020).

### Software

We employed customized scripts in python (version 3.8.11), using matplotlib (v3.6.0), seaborn (v0.11.1), and biopython (v1.78) libraries.

## Supplementary Figures

**Supplementary Figure 1:** A) Bar plot comparing the performance of different model architectures (MLP, XGBoost, KRR and SMRF) and input feature representations (cell features and drug features) in terms of Pearson correlation with observed drug sensitivities. Hatched bars highlight models using only trascriptomics data; B) Bar plot depicting Mean Squared Error (MSE) for the same models and features as in (A);

**Supplementary Figure 2:** Evaluation of PRISM models across target protein families and cell line responses. A) Bar chart depicting the number of drugs targeting specific protein families. Light blue bars represent the total count of drugs, while dark blue bars indicate the count of drugs with an interquartile range (IQR) greater than 1, highlighting the variability in response within each protein family; B) Bar chart of models stratified by putative target protein families with a correlation coefficient (corr) greater than 0.2. Salmon bars represent the models with corr > 0.2, while dark red bars indicate models with corr > 0.2 and at least one target identified; C) Boxplot distribution of the number of drugs per protein family showing negative fold changes (FC) in cell viability at thresholds of −1.5FC, −2FC, and −3FC, in relation to the median FC. This visualizes the sensitivity of cell lines to drugs within each target family, with the accompanying bar chart below summarizing the total count of drugs per family that induce a FC of at least −1.5.

**Supplementary Figure 3:** Figure A) Heatmap of LLM Pathway Recovery Probability by Drug Category. This heatmap illustrates the likelihood of L1 pathways being identified as relevant within various drug categories. Each row represents one of the top 25 most frequently detected pathways, while columns correspond to different drug categories. The color gradient, ranging from yellow (low probability) to blue (high probability), represents the ratio of occurrences of a given pathway within a drug category to the total number of drugs in that category. Values are represented numerically within the cells. The plot shows how the LLM pathway selection procedure delivers qualitatively different pathways across different drug categories. B) Venn diagram representing the number of unique pathways shared by chemotherapeutics, targeted drugs and non-cancer drugs.

**Supplementary Figure 4:** A) boxplot of correlation distributions for GDSC models showing the greatest correlation worsening of the MOA-primed vs all-genes models; B) boxplot of correlation distributions for PRISM models showing the greatest correlation worsening of the MOA-primed vs all-genes models

**Supplementary Figure 5:** A) mutational burden of oncodrivers in top 1000 patients scored by the Dabrafenib model; B) number of samples, among the top 1000 scored by Dabrafenib, with BRAF mutations grouped by cancer type.

**Supplementary Figure 6:** stacked barplot of the GDSC drugs scoring among the top 1000 samples patients with cancer types matching the prescription according to NCI drugs. The height of the barplot’s stacks corresponds to relative count of unique samples and the color of the specific cancer type. TCGA codes marked with the asterisk “*” highlight tumor types for which the drug is prescribed (source NCI).

**Supplementary Figure 7:** CellHit predictions on distinct PDAC subtypes. A) UMAP plot of cancer cell lines and the PDAC subtypes (average of samples annotated according to their subtype: GL, Glandular; TR, Transitional; UN, Undifferentiated) after their integration by Celligner. Top left: dashed boxes indicate the space of three PDAC subtypes shown at higher magnification on the top. Top right: pie chart showing the percentage of predicted cancer cell lines that react most similarly to the available drugs for all PDAC samples. Classification of the tissue types of cell lines (oncotree system) is shown.

**Supplementary Figure 8:** clustering of CellHit predictions of GDSC drugs (rows) for each GBM sample (columns). Cells contain the predicted lnIC50 values normalized by median subtraction. Columns are color annotated via sample metadata, including tumor type, i.e. primary (purple) or recurrent (yellow), and response type, i.e. non-response (red) or response (green). Vertical green rectangles indicate tested samples, horizontal green rectangles indicate tested drugs.

## Supplementary Tables

**Supplementary Table S1: comparative performance of drug-response prediction models trained on GDSC data**

**Supplementary Table S2: GDSC drug individual models performances and important genes**

**Supplementary Table S3: reactome pathways associated to GDSC’s drugs mechanisms of actions**

**Supplementary Table S4: Reactome pathway enrichment on GDSC’s models important genes**

**Supplementary Table S5: GDSC’s models important genes (based on SHAP) recall of essential genes across cancer tissues**

**Supplementary Table S6: GDSC’s models SHAP importances Supplementary Table S7: PRISM drug individual models performances**

**Supplementary Table S8: PRISM drug individual models performances (corr > 0.2) and important genes for models with known target**

**Supplementary Table S9: reactome pathways associated to PRISM’s drug mechanisms of actions**

**Supplementary Table S10: Reactome pathway enrichment on PRISM’s models important genes**

**Supplementary Table S11: list of FDA approved drugs for a specific cancer type**

**Supplementary Table S12: TCGA sample inference output and drug prescription match information**

**Supplementary Table S13: predicted drug combinations and support**

**Supplementary Table S14: approved drug combinations from Drugbank**

**Supplementary Table S15: CellHit inference on PDAC patient samples**

**Supplementary Table S16: CellHit inference on GBM patient samples**

## Data availability

The generated dataset are available at:

https://cellhit.bioinfolab.sns.it

## Code availability

The code of the model and pipelines employed in this study are available at:

https://github.com/raimondilab/CellHit

## Acknowledgments

F.R. was supported by the Italian Ministry of University and Research through the Department of Excellence “Faculty of Sciences” of Scuola Normale Superiore. The research leading to these results also received funding from the Italian Association for Cancer Research (AIRC) under My First AIRC Grant (MFAG) 2020 - ID. 24317 project. We gratefully acknowledge the CINECA award, in collaboration with AIRC, for the availability of high-performance computing resources and generous support. We gratefully acknowledge the computational resources of the Center for High-Performance Computing (CHPC) at Scuola Normale Superiore. G.N. was supported by an AIRC Investigator Grant #20251 and AIRC 5×1000 Grant #21147. This work was also partially supported by the Italian Ministry of Health with the “Ricerca Corrente” and “5×1000” funds to the IEO IRCCS.

## Notes

### Competing Interest Statement

The authors have declared no competing interest.

### Summary of Updates

Author list updated. Supplemental files updated

